# A thermodynamic atlas of carbon redox chemical space

**DOI:** 10.1101/245811

**Authors:** Adrian Jinich, Benjamin Sanchez-Lengeling, Haniu Ren, Joshua E. Goldford, Elad Noor, Jacob N. Sanders, Daniel Segrè, Alán Aspuru-Guzik

## Abstract

Redox biochemistry plays a key role in the transduction of chemical energy in living systems. However, the compounds observed in metabolic redox reactions are a minuscule fraction of chemical space. It is not clear whether compounds that ended up being selected as metabolites display specific properties that distinguish them from non-biological compounds. Here we introduce a systematic approach for comparing the chemical space of all possible redox states of linear-chain carbon molecules to the corresponding metabolites that appear in biology. Using cheminformatics and quantum chemistry, we analyze the physicochemical and thermodynamic properties of the biological and non-biological compounds. We find that, among all compounds, aldose sugars have the highest possible number of redox connections to other molecules. Metabolites are enriched in carboxylic acid functional groups and depleted of carbonyls, and have higher solubility than non-biological compounds. Upon constructing the energy landscape for the full chemical space as a function of pH and electron donor potential, we find that over a large range of conditions metabolites tend to have lower Gibbs energies than non-biological molecules. Finally, we generate Pourbaix phase diagrams that serve as a thermodynamic atlas to indicate which compounds are local and global energy minima in redox chemical space across a set of pH values and electron donor potentials. Our work yields insight into the physicochemical principles governing redox metabolism, and suggests that thermodynamic stability in aqueous environments may have played an important role in early metabolic processes.

## Introduction

Redox reactions are fundamental to biochemistry. The two main biogeochemical carbon-based transformations - respiration and photosynthesis - are at heart oxidative and reductive processes, and a large fraction of catalogued enzymatic reactions (≈ 40%) are oxidoreductive in nature ^1,2^. Thermodynamics and other physicochemical properties act as constraints on the evolution of metabolism in general and of redox biochemistry in particular. A classic example is the adaptation and expansion of metabolism in response to Earth’s great oxidation event (GOE) ^3–6^. The rise in molecular oxygen resulted in a standard redox potential difference of ≈ 1.1 eV available from NAD(P)H oxidation, and led to the emergence of novel biochemical pathways such as the biosynthesis of sterols ^7–9^.

Recent work has uncovered quantitative thermodynamic principles that influence the evolution of carbon redox biochemistry ^10–12^. This line of work has focused on the three main types of redox reactions that change the oxidation level of carbon atoms in molecules: reductions of carboxylic acids (-COO) to carbonyls (-C=O); reductions of carbonyls to alcohols (hydroxycarbons) (C-O), and reductions of alcohols to hydrocarbons (C-C). The “rich-get-richer” principle states that more reduced carbon functional groups have higher standard redox potentials ^10–12^. Thus, alcohol reduction to a hydrocarbon is more favorable than carbonyl reduction to an alcohol, which in turn is more favorable than carboxylic acid reduction to a carbonyl. This explains why, across all six known carbon fixation pathways, ATP is invested solely (with Ribulose-5P kinase as the single exception) to drive carboxylation and the reduction of carboxylic acid functional groups ^11,13^. Quantitative analysis of biochemical redox thermodynamics has also explained the emergence of NAD(P) as the universal redox cofactor. With a standard redox potential of −320 mV, NAD(P) is optimized to reversibly reduce/oxidize the vast of majority of central metabolic redox substrates ^12^. In addition, since its standard potential is approximately 100 mV lower than that of the typical carbonyl functional group, it effectively decreases the steady-state concentration of potentially damaging carbonyls in the cell^12^. Finally, other physicochemical properties like hydrophobicity and charge act as constraints that shape the evolution of metabolite concentrations ^14^.

Despite this knowledge, the thermodynamic and physicochemical principles underlying the rise of carbon redox biochemistry remain very poorly understood. Here we combinatorially generate the chemical space of all possible redox states of linear-chain n-carbon compounds (for n=2-5). We partition each n-carbon linear-chain redox chemical space into biological metabolites and non-biological compounds, and systematically explore whether metabolites involved in biochemical redox reactions display features that would be unexpected elsewhere in redox chemical space. To compare physicochemical and thermodynamic properties of the biological and non-biological molecules we use cheminformatic tools and a recently developed quantum chemical approach to estimate standard reduction potentials (E^o^’)^12^ of biochemical reactions. In addition to generating a molecular energy landscape of broad applicability to the study of biochemical evolution, our analysis provides specific insight on redox biochemistry. In particular, we find that (1) the oxidation level and asymmetry of aldose sugars makes them unique in that they have the highest possible number of connections (reductions and oxidations) to other molecules; (2) biological compounds (metabolites) tend to be enriched for carboxylic acid functional groups and depleted for carbonyls; (3) metabolites tend to have, on average, higher solubilities and lower lipophilicities than the non-biological molecules; (4) across a range of pH and electron donor/acceptor potentials metabolites tend to have, on average, lower Gibbs energies relative to the non-biological compounds; (5) by adapting Pourbaix phase diagrams - an important conceptual tool in electrochemistry - to the study of redox biochemistry, we find that the n-carbon linear-chain dicarboxylic acids and fatty acids (e.g. succinate and butyrate in 4-carbon redox chemical space) are the local minima in the energy landscape across a range of conditions, and thus may have a spontaneous tendency to accumulate. Our results suggest that thermodynamics may have played an important role in driving the rise of dominant metabolites at the early stages of life, and yields insight into the principles governing the emergence of metabolic redox biochemistry.

## Results

### Aldose sugars have the maximal number of redox connections

We combinatorially generated all possible redox states of n-carbon linear-chain compounds (for n = 2-5 carbon atoms per molecule) and studied the properties of the resulting chemical spaces (Fig. 1). For every molecule in n-carbon redox chemical space, each carbon atom can be in one of four different oxidation levels: carboxylic acid, carbonyl (ketone or aldehyde), hydroxycarbon (alcohol), or hydrocarbon (Fig. 1A). Molecules in redox chemical space are connected to each other by three different types of 2-electron reductions (or the reverse oxidations) that change the oxidation level of a single carbon atom: reduction of a carboxylic acid to a carbonyl; reduction of a carbonyl to a hydroxycarbon; and reduction of a hydroxycarbon to a hydrocarbon. In order to make the redox chemical space model system tractable to analysis, we decreased its complexity by excluding carbon-carbon bond cleavage/formation reactions (e.g. reductive carboxylations or oxidative decarboxylations), keto-enol tautomerizations, intermediate carbon-carbon double-bond formation, intramolecular redox reactions, or different stereoisomers for a given molecular oxidation level. In what follows, we focus the majority of our analysis on the properties of the 4-carbon linear-chain redox chemical space. (See SI Figures 7 - 14 for corresponding results in 2-, 3-, and 5-carbon linear-chain redox chemical space).

**Figure 1.**
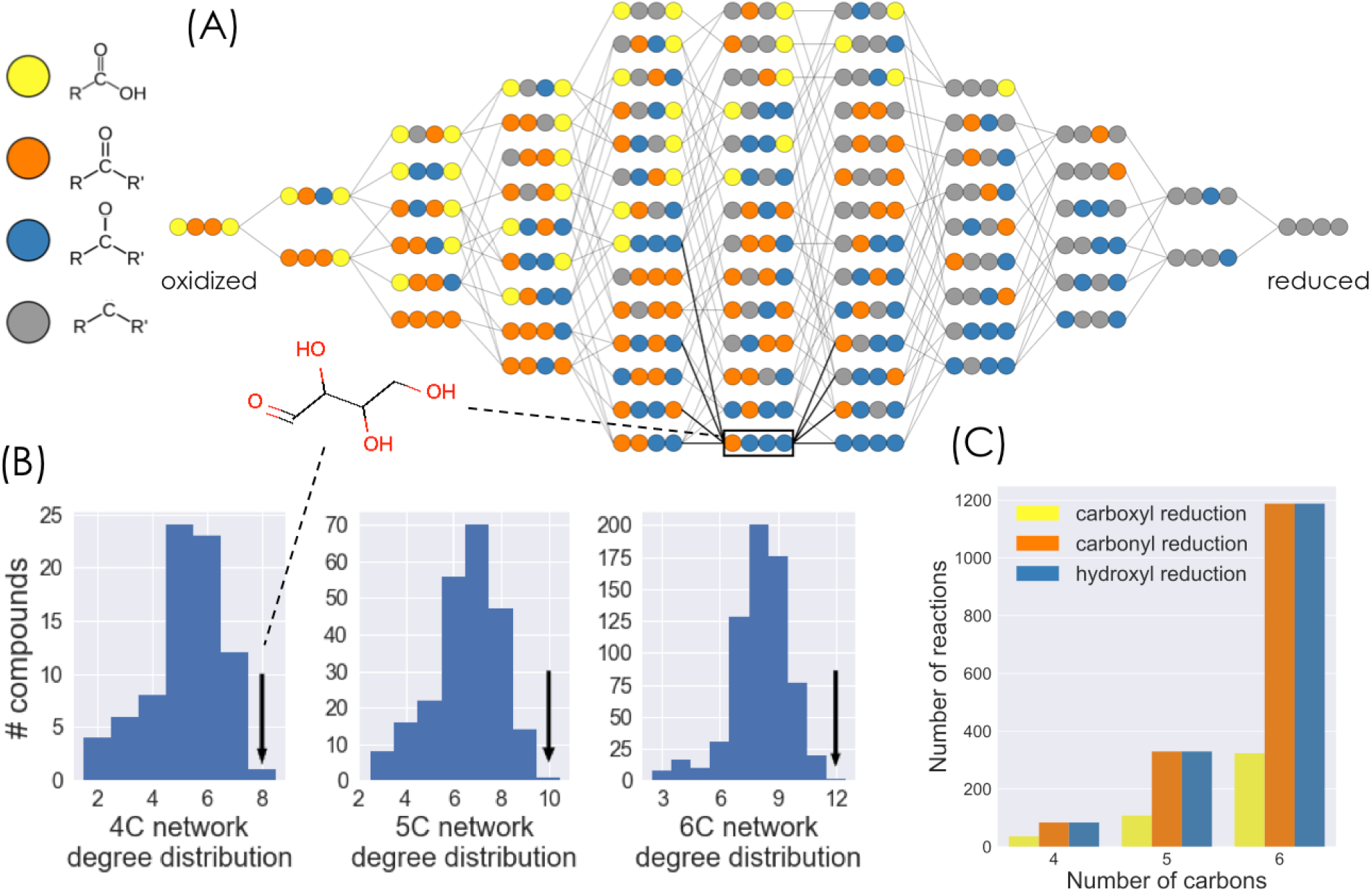
The structure of n-carbon linear-chain redox chemical space. A) The redox chemical space defined by the set of all possible 4-carbon linear chain molecules that can be generated from three different types of redox reactions: reduction of a carboxylic acid to carbonyl group; reduction of a carbonyl group to a hydroxycarbon (alcohol); and reduction of a hydroxycarbon to a hydrocarbon (and corresponding oxidations). Carbon atoms are represented as colored circles, with each color corresponding to an oxidation state: yellow = carboxylic acid; orange = carbonyl; blue = hydroxycarbon; gray = hydrocarbon. Compounds within each column have the same molecular oxidation state and are organized from most oxidized (left) to most reduced (right). B) The degree distributions for the 4-, 5-, and 6-carbon linear chain redox chemical spaces. In all cases, the aldose sugar is the only compound with the maximal number of possible reductions and oxidations (black arrows). C) Number of reactions in the 4-, 5-, and 6-carbon linear chain redox chemical spaces that belong to each of the three types of redox reactions considered.

The 4-carbon linear-chain redox chemical space contains 78 molecules connected by 204 reactions. The molecules span 11 different molecular oxidation levels, from the fully oxidized 2,3-dioxosuccinic acid (two carboxylic acids and two carbonyls) to the fully reduced alkane butane (Figure 1a). 84 reactions reduce carbonyls to hydroxycarbons (or oxidize hydroxycarbons to carbonyls), and the same number reduce hydroxycarbons to hydrocarbons (or oxidize hydrocarbons to hydroxycarbons). Since carboxylic acids are restricted to carbon atoms at the edges of a molecule (i.e. carbons #1 and #4 in 4-carbon linear-chain molecules), only 36 reactions reduce carboxylic acids to aldehydes (or oxidize aldehydes to carboxylic acids) (Fig 1c).

The number of reactions that connect a molecule to its oxidized or reduced products - the redox degree of a molecule - ranges from 2 to 2n (Fig 1b). In n=4-carbon redox chemical space, we find that only a single molecule in the network, the aldose sugar erythrose (and its stereoisomers), has the maximal degree value of 2n=8. This holds true for all redox chemical spaces regardless of the number of carbon atoms: only the corresponding aldose sugars in the 2-, 3-, 5-, and 6-carbon redox chemical spaces have the maximal degree value, 2n (Fig 1B). This is explained by the fact that the n-carbon aldose sugar satisfies the two constraints required to have the maximal number of redox connections: (i) each atom must be in an “intermediate” oxidation level that can be both oxidized and reduced. Therefore all inner carbon atoms (i.e. atoms #2 and #3 in 4-carbon linear-chain molecules) must be in the hydroxycarbon oxidation level, while carbon atoms at the edges (i.e. atoms #1 and #4) can be either in the carbonyl (aldehyde) or hydroxycarbon oxidation level. (ii) The molecule must not be symmetric under a 180 degree rotation along its center. Thus the two edge atoms must be in different oxidation levels. This leads uniquely to the aldose sugar molecular redox state configuration.

### Biological compounds are enriched in carboxylic acids and depleted of carbonyl groups

What distinguishes the subset of compounds in redox chemical space that appear in cellular metabolism from those that do not? To address this question we subdivided the 78 molecules from the full 4-carbon redox chemical space into 30 biological compounds (also referred to from here onwards simply as metabolites or “natural” compounds), which were identified based on matches with KEGG database entries ^1,2^, and the remaining 48 “non-biological” compounds (Fig. 2A). Compounds in KEGG that correspond to molecules in redox chemical space but have alcohol groups substituted by amines or phosphates were considered a match, as these functional groups have the same oxidation level (see Methods for further details). For example, the metabolites oxaloacetate and aspartate have the same oxidation level at every carbon atom but differ by the substitution of an alcohol into an amine; both are considered a match to the corresponding molecule in our network. Similarly, we consider metabolites with carboxylic acid groups that are activated with either thioesters or phosphates groups as matches to molecules in redox chemical space.

**Figure 2.**
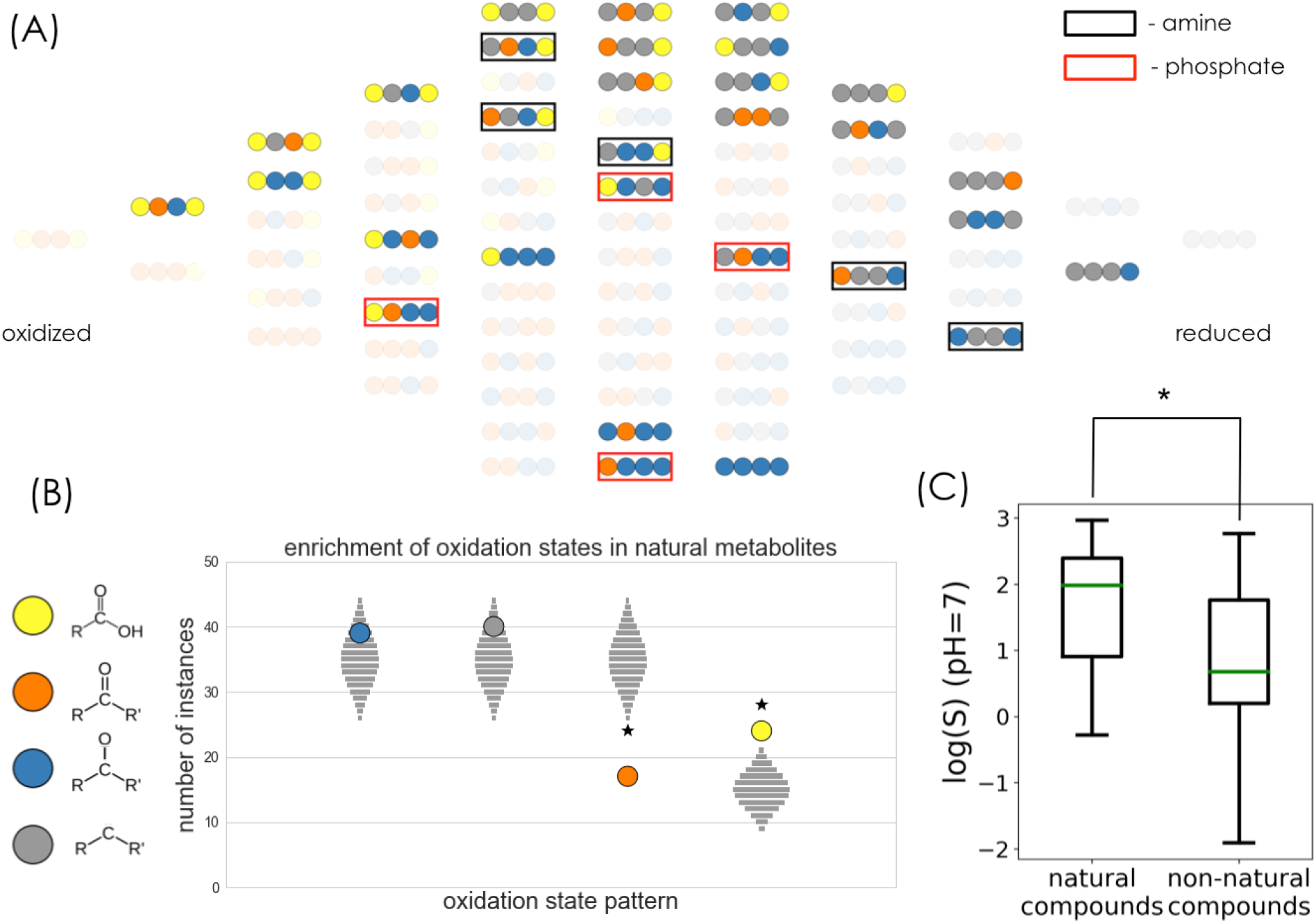
Functional group statistics and aqueous solubilities of biological compounds in the 4-carbon linear chain redox chemical space. A) The subset of molecules in 4-carbon linear-chain redox chemical space that match biological metabolites in the KEGG database. Compounds that match KEGG metabolites but with alcohol groups substituted by either amines or phosphates are marked with black and red squares, respectively. B) Enrichment and depletion of functional groups in the set of biological compounds. The vertical position of each colored circle corresponds to the number of times each functional group appears in the set of biological compounds. The light gray squares show the corresponding expected null distributions for random sets of molecules sampled from redox chemical space. See Fig S1 for statistical analysis of functional group pairs and triplets. C) Comparison of predicted aqueous solubility log(S) at pH=7 for biological and non-biological compounds in the 4-carbon linear-chain redox chemical space. biological compounds have significantly higher solubilities than the non-biological set (p < 0.005).

As a first comparison between metabolites and non-biological compounds, we analyzed the enrichment or depletion of functional groups (i.e. carbon atom oxidation levels) in the two categories (Fig. 2B). Specifically, we counted the number of times that each functional group appears in the set of metabolites, and compared it to analytically-derived expected null distributions for random sets of compounds (see Methods). We found that in 4-carbon linear-chain redox chemical space, metabolites are significantly enriched in carboxylic acids (p<0.001) while being significantly depleted for ketones (p<0.001) (Fig. 2B). We find similar trends in 3- and 5-carbon redox chemical spaces (Figs. S8 and S11. Since all but one molecule in 2-carbon redox chemical space are biological metabolites, this space is not amenable to such statistical analysis). Furthermore, after normalizing for observed single functional group statistics (see Methods for further details) we computed the null distributions for higher order functional group patterns, i.e. pair (2-mer) and triplet (3-mer) patterns (Fig. S1). According to our analysis, only the 2-mer pattern with a hydroxycarbon next to a hydrocarbon is depleted in the metabolites, albeit not significantly (p=0.05). The number of times that all other 2-mer and 3-mer functional group patterns appear in metabolites - including the highly uncommon dicarbonyl pattern - can be explained by the underlying single functional group statistics.

We then asked whether the observed functional group enrichments and depletions translate to differences in physicochemical properties of the metabolites and the non-biological compounds. Towards this end, we used cheminformatic tools to estimate the values of solubility (logS) and lipophilicity (as captured by the octanol-water partition coefficient, logP) at pH=7. We find that, in correlation with their enrichment for carboxylic acid functional groups, metabolites have significantly higher solubilities (p<0.005) (Fig 2C), and significantly lower octanol-water partition coefficients (p<0.01) (Fig S2) than the set of non-biological compounds. We observe similar trends in 3- and 5-carbon redox chemical spaces (Figs. S9 and S13).

### Metabolites have on average lower Gibbs energies than non-biological compounds

We next focused on estimating the energy landscape of our redox compounds, with special attention to the question of whether metabolites and non-biological compounds display different patterns in this landscape. We used a recently developed calibrated quantum chemistry approach ^12^ to accurately predict the apparent standard redox potentials *E*^*o*^′(*pH*) of all reactions in n-carbon linear-chain redox chemical space (n = 2-5). Previous work has shown that the calibrated quantum chemistry method achieves significantly better accuracy than group contribution method (GCM)^12^, the most commonly used approach to estimate thermodynamic parameters of biochemical compounds and reactions ^15–18^. Briefly, the quantum chemistry method relies on density functional theory (DFT) with a double-hybrid functional ^19,20^ to compute the differences in molecular electronic energies and utilizes a two-parameter calibration against available experimental data. We computed the energies of several geometry-optimized conformations of the fully protonated species of each compound. We then estimated the standard redox potential *E*^*o*^ of the fully protonated species as the difference in electronic energies of the products and substrates, ΔE_electronic_. Using cheminformatic pKa estimates (Marvin 17.7.0, 2017, ChemAxon) and the Alberty Legendre transform ^21,22^, we converted the standard redox potentials to *transformed* standard redox potentials *E*^*o*^′(*pH*), which depend on pH. Finally, in order to correct for systematic errors in the quantum chemistry calculations and the cheminformatic pKa estimates, we calibrated - via linear regression - the transformed standard redox potentials *E*^*o*^′(*pH*) against a dataset of available experimental values (see Methods for further details).

We note that the improvement in accuracy of the quantum chemical approach over GCM is particularly striking for the linear-chain compounds in our redox chemical spaces. This is most apparent for the set of carbonyl to hydroxycarbon reductions (Fig S3): while GCM prediction is no better than an average value predictor (R^2^=−0.04), the redox potentials predicted with the calibrated quantum chemistry method correlate linearly with experimental values (Pearson r=0.50). GCM accounts only for the difference in group energies of products and substrates to estimate redox potentials. Thus for redox reactions it effectively ignores the molecular environment surrounding the reduced/oxidized carbon atom, collapsing all the potentials associated to carbonyl functional group reductions to two values (the average aldehyde and ketone reduction energies) thus lowering its prediction accuracy (Fig S3). Therefore the use of our calibrated quantum chemical method is essential in order to accurately predict and analyze the energetics of n-carbon linear chain redox chemical spaces.

We used the predicted *E*^*o*^′(*pH*) values to generate the energy landscape of redox chemical space. To do this, we assumed that each compound is coupled to an electron donor/acceptor with a given steady-state redox potential, *E*(*electron donor*). This potential could represent that set by a steady state ratio of NAD^+^/NADH or other abundant redox cofactor inside the cell^23^. Alternatively, in the context of prebiotic chemistry, it could represent the potential associated to a given concentration of molecular hydrogen in an alkaline hydrothermal vent or different iron oxidation states in prebiotic oceans ^24,25^. Given a value of *E*(*electron donor*), we convert the *E*^*o*^′(*pH*) of each reaction into a Gibbs reaction energy, using Δ*G*_r_ (*pH*) = − *nF* (*E*^*o*^′(*pH*) − *E*(*electron donor*)) (where n is the number of electrons and F is Faraday’s constant). The set of Gibbs reaction energies for all redox transformations - as a function of pH and electron donor potential - defines the energy landscape of our redox chemical space (Fig. 3A, S5, S6)

**Figure 3.**
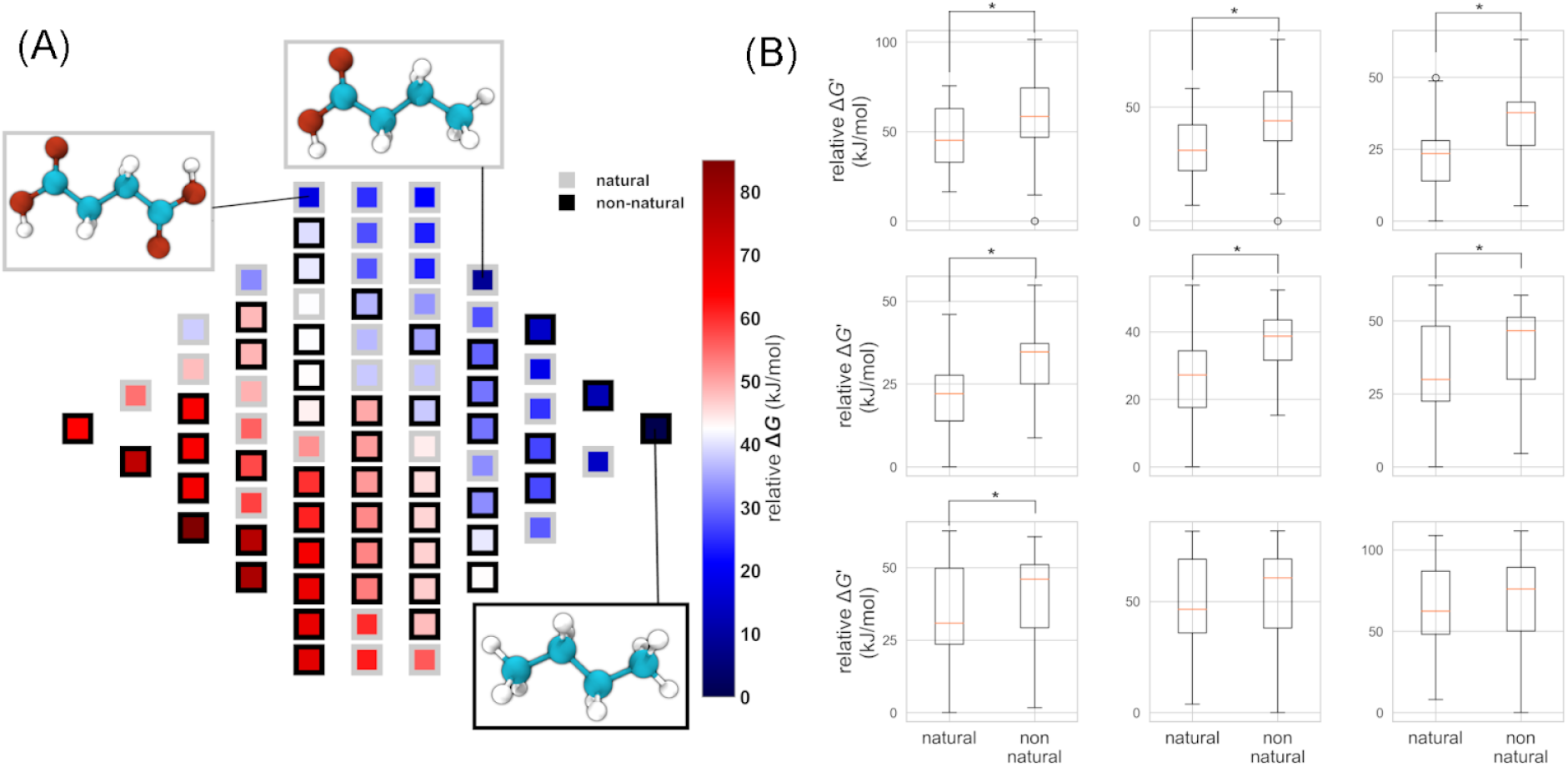
Thermodynamic landscape of the 4-carbon linear chain redox chemical space. A) Relative Gibbs energies of metabolites at pH=7 and E(electron donor/acceptor) = −300 mV. Gibbs energies are normalized relative to the metabolite with the lowest energy. Compounds within a column (same molecular oxidation state) are sorted from highest (bottom) to lowest (top) relative energies. The structures of the three compounds that are local minima in the thermodynamic landscape are shown: succinate (left), butyrate (top), and butane (bottom right). These compounds have lower Gibbs energies than all their neighboring molecules accessible by a reduction or oxidation. B) Relative Gibbs energies of biological and non-biological compounds for a range of pH and E(electron donor/acceptor) values. At each value of pH and E(electron donor/acceptor), Gibbs energies are normalized relative to the compound with the lowest energy. Asterisks indicates statistically significant differences of average values (Welch’s t-test, p<0.05).

A notable finding of this analysis is that, across a range of cofactor potentials, metabolites in 4-carbon linear-chain redox chemical space have on average significantly lower relative Gibbs energies than the non-biological compounds (Fig 3B). We find a similar trend for metabolites in 3- and 5-carbon linear-chain redox chemical space (Fig. S10, S14); however because of the few number of non-natural compounds in 3-carbon redox chemical space, the trend there is not statistically significant (Fig. S10). An important exception to this general trend (and one that is conserved across spaces with different numbers of carbon atoms) is that the aldose sugars (e.g. erythrose), the ketose sugars (e.g. erythrulose), and the sugar alcohols (e.g. threitol) have a higher relative Gibbs energy than all compounds in redox chemical space across a large range of pH and electron donor potential.

### A Pourbaix phase diagram of redox chemical space maps local minimal energy compounds

In addition to the trends observed for average energy differences between biological and non-biological compounds, the relative energies of individual compounds change as a function of pH and *E*(*electron donor*)(Fig S5, S6). To further investigate the detailed structure of the thermodynamic landscape, we set out to map which molecules are local minima at each value of pH and *E*(*electron donor*). A molecule is a local minimum in redox chemical space if its Gibbs energy is lower than that of all its neighbors with whom it is connected through a reduction or an oxidation. We adapt Pourbaix phase diagrams, a powerful standard visualization tool in the field of electrochemistry ^26^, to the problem of mapping out regions of pH-*E*(*electron donor*) phase space in n-carbon linear-chain redox chemical space. In a Pourbaix diagram, the predominant equilibrium states of an electrochemical system and the boundaries between these states are mapped out as a function of two phase space parameters. Fig. 4 shows a Pourbaix phase diagram representation of 4-carbon linear-chain redox chemical space. (See Fig. S7, S8, and S12 for the corresponding Pourbaix diagrams for 2-, 3-, and 5-carbon linear-chain redox chemical spaces).

**Figure 4.**
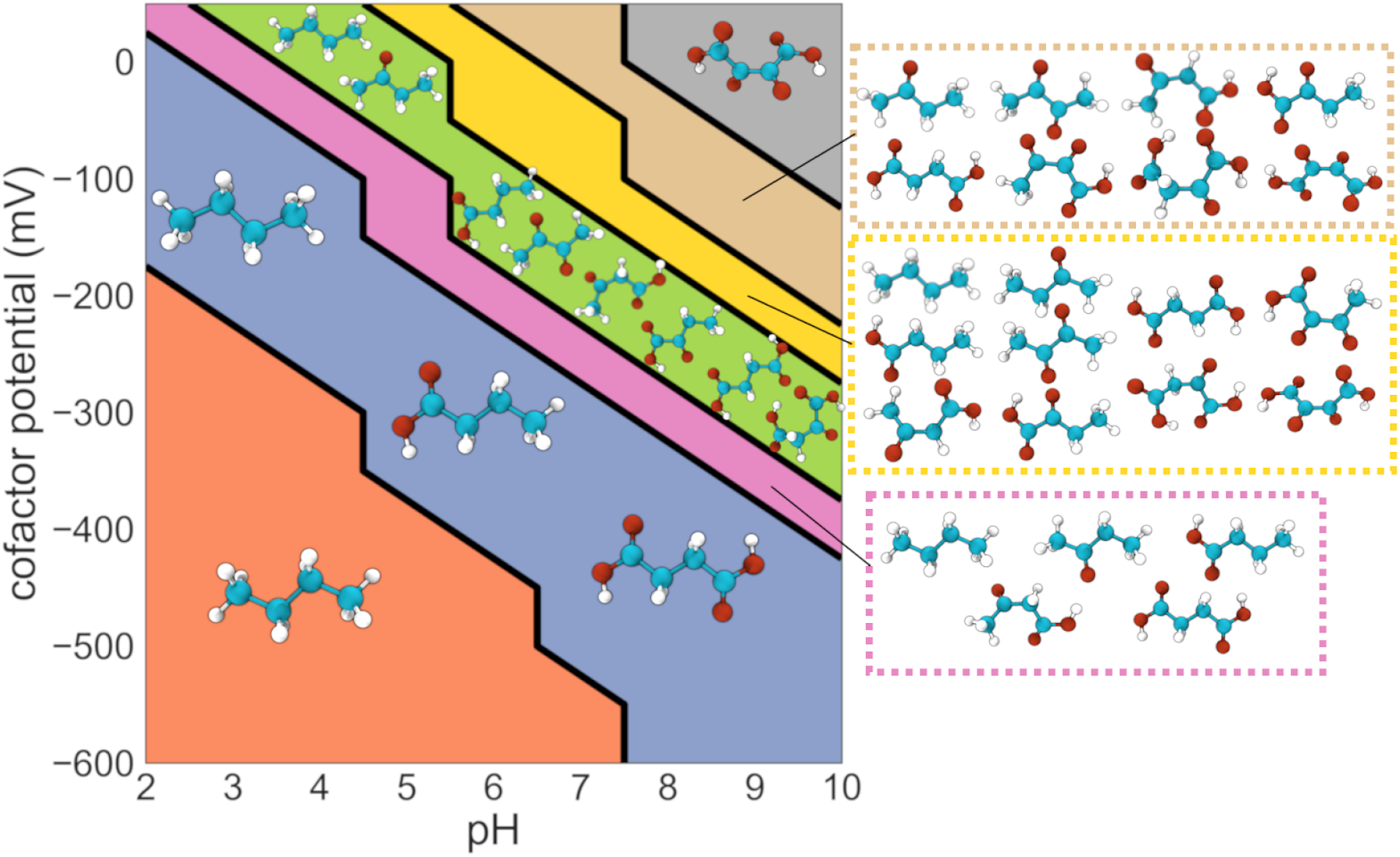
Pourbaix phase diagram for the 4-carbon linear chain redox chemical space. Molecules that are local minima in the energy landscape at each region of pH, E(electron donor/acceptor) phase space are shown. At low pH and E(electron donor/acceptor) values, butane is both the global and the only local minimum energy compound. At intermediate values of pH and E(electron donor/acceptor), several metabolites emerge as local minima and would thus tend to accumulate. For example, the metabolites oxaloacetate, acetoacetate, and alpha-ketobutyrate emerge as local energetic minima in the region of phase space shown in green. Finally, in the upper right corner of the phase diagram, characterized by higher values of both pH and E(electron donor), the fully oxidized four carbon compound 2,3-dioxosuccinic acid emerges as the only local (global) minimum.

At the lower left corner of the diagram, in the region corresponding to more acidic pH values and more negative electron donor potentials, the fully reduced 4-carbon alkane butane is the only local (and the global) energy minimum (Fig. 4). Thus, assuming all compounds are kinetically accessible, butane would be expected to accumulate in these conditions. The structure of redox chemical space becomes richer as pH and *E*(*electron donor*) increase. Succinate and the 4-carbon short-chain fatty acid (SCFT) butyrate - two biologically important metabolites - emerge as two additional local minima at more oxidative regions of the phase diagram (Fig. 4). Both succinate and butyrate consist of inner carbon atoms in the hydrocarbon (fully reduced) oxidation level, and edge carbon atoms in either the hydrocarbon or the carboxylic acid (fully oxidized) state. Notably this pattern - where the n-carbon linear-chain dicarboxylic acid (oxalate, malonate, succinate, and glutarate for 2-, 3-,4-, and 5-carbon atoms, respectively), the fatty acid (acetate, propionate, butyrate, and valerate), and the alkane (ethane, propane, butane, and pentane) emerge as the only local minima in a large region of phase space - is conserved in redox chemical spaces with different number of carbon atoms (Fig. S7, S8, and S12). Further increases in either pH or electron donor potential result in the emergence of additional compounds, both metabolites and non-biological molecules, as local energy minima in the landscape (Fig. 4). (See Fig. S7, S8, S12).

Can we predict from simple physicochemical principles the identity of the local minimal energy compounds? A simple mean-field toy model (Fig. 4c) that focuses on the average standard redox potentials < *E*^*o*^′(*pH*) > of the different carbon functional groups can help intuitively predict which metabolites accumulate at given values of pH and *E*(*electron donor*). Fig. 5 (upper panel) shows the distributions of standard potentials at a fixed pH (pH=7) for all compounds in 4-carbon redox chemical space categorized by the type of functional group undergoing reduction. Given a fixed value of *E*(*electron donor*), the average redox potentials for each functional group category can be used to compute average Gibbs reaction energies for each type of carbon redox transformation via the following equation: Δ*G*_r_ (*pH*) = − *nF* (< *E*^*o*^′(*pH*) > − *E*(*electron donor*)) (where n is the number of electrons and F is Faraday’s constant). We use this to generate Fig. 5 (lower panel), which schematically shows the relative average Gibbs energies of the four different carbon oxidation levels at different values of *E(electron donor)* at pH=7. The boundaries delimiting different regions of the *E(electron donor)* axis mark the values where the rank-ordering of relative average Gibbs energies for the four carbon oxidation levels changes.

**Figure 5.**
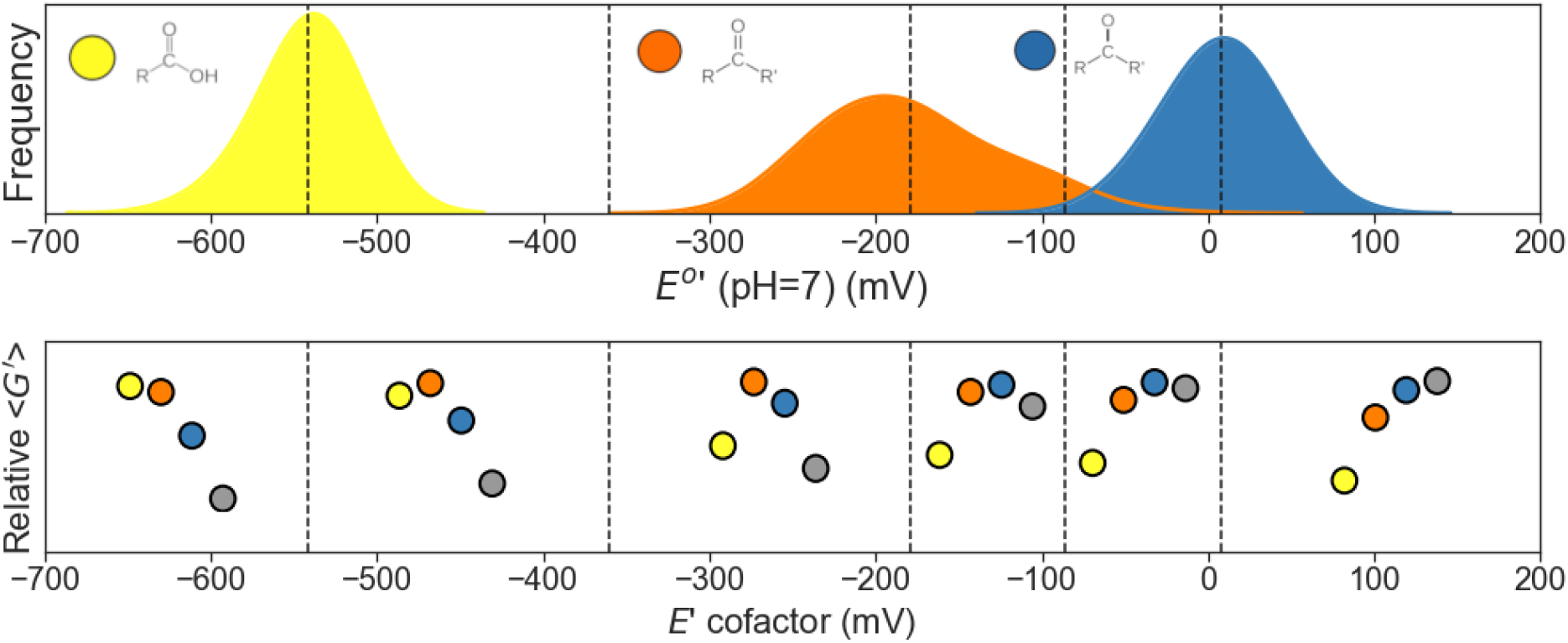
A mean-field toy model explains the identity of molecular oxidation states of local minima. The schematic diagram illustrates how the average standard redox potentials of different carbon functional groups dictate the identity of the minimal energy compounds. The top panel shows the distributions of standard redox potentials (pH=7) of all reactions in the 4-carbon linear-chain redox chemical space, grouped according to the functional group that is reduced during the transformation: carboxylic acid (yellow), carbonyl (orange), and hydroxycarbon (blue). The bottom panel shows - for different values of E(electron donor/acceptor) - the resulting relative average Gibbs energies of the functional groups. For example, in the region where E(electron donor/acceptor) is between about −360 and −190 mV, carbonyls (orange) have on average the highest relative Gibbs energy, followed by hydroxycarbons (blue), carboxylic acids (yellow), and hydrocarbons (gray). Therefore minimal energy compounds will have *inner* carbon atoms (atoms #2 and #3 in 4-carbon molecules) that equilibrate to the hydrocarbon oxidation state, and *edge* carbon atoms (atoms #1 and #4) that equilibrate to either the carboxylic acid or the hydrocarbon oxidation state.

As an illustrative example, we focus on region III in Fig. 5, which is approximately delimited by the values − 360 *mV* ≤ *E*(*electron donor*) ≤ − 180 *mV*. In this region, the reduction of carboxylic acids to aldehydes (yellow to orange) is highly unfavorable (alternatively, the oxidation of aldehydes to carboxylic acids is highly favorable). On the other hand, the reduction of ketones to hydroxycarbons (orange to blue), as well as the reduction of hydroxycarbons to hydrocarbons (blue to gray) are thermodynamically favorable. Thus at pH=7 and for this range of *E(electron donor)* values, the edge carbon atoms (atoms #1 and #4 in the 4-carbon linear-chain compounds) are driven to either the most oxidized (carboxylic acid) or the most reduced (hydrocarbon) oxidation level, while the inner carbon atoms (atoms #2 and #3) - which cannot exist in the carboxylic acid oxidation level - are driven to the hydrocarbon oxidation level. This corresponds precisely to the molecular oxidation levels of dicarboxylic acids, fatty acids, and alkanes (e.g. succinate, butyrate, and butane), the local and global minimal energy compounds in this region of pH-E(electron donor) phase space.

## Discussion

In this work, we introduced the chemical spaces of all molecular oxidation levels of n-carbon linear-chain compounds, and analyzed their structural, physicochemical, and thermodynamic properties. Examining the connectivity of redox chemical space, we found that aldose sugars - e.g. glyceraldehyde (n=3), erythrose (n=4), ribose (n=5), and glucose (n=6), and their corresponding stereoisomers - are unique in that they are the only compounds with the highest possible number of oxidative and reductive connections (2n) to neighboring molecules. Whether this maximal number of connections played a role in the emergence of aldose sugars as key players in cellular metabolism remains to be explored.

We found that the set of biological compounds is significantly enriched for carboxylic acid functional groups and have, on average, significantly higher solubilities (logS at pH=7) than the set of non-biological compounds. In addition to an increase in aqueous solubility, other reasons why carboxylic acids may have been selected during the evolution of metabolism potentially include a decrease in permeability across lipid membranes ^27^. This is reflected in the predicted values of octanol-water partition coefficients, logD(pH=7) for the biological and non-biological compounds (Fig. S2). In addition, the enrichment for carboxylates may have enhanced the ability of enzymes to recognize small molecule substrates. Our analysis also showed that biological compounds are significantly depleted in carbonyl functional groups. Notably, in the 4-carbon network only one biological compound, diacetyl - which appears in the metabolic networks of yeast and several bacterial species ^28,29^ - contains two carbonyl functional groups. This is consistent with the fact that carbonyl groups are significantly more reactive than carboxylic acids or hydroxycarbons, and can cause oxidative damage, spontaneously cross-link proteins, inactivate enzymes and mutagenize DNA ^30^.

Our thermodynamic calculations, which rely on a recently developed calibrated quantum chemistry approach ^12^, revealed that metabolites have on average lower Gibbs energies than the non-biological set of compounds across a range of pH and electron donor potentials. This finding provides quantitative evidence for the reasonable yet “highly speculative” notion put forward by Bloch and others that, during life’s origins, “the thermodynamically most stable compounds had the best chance to accumulate and survive” ^31^.

The resulting thermodynamic landscape also revealed in detail which compounds - both biological and non-biological - are energetic local minima as a function of pH and *E(electron donor)*, and would thus tend to accumulate. For example, as captured in the Pourbaix phase diagram representation of 4-carbon linear-chain redox chemical space, the biological metabolites succinate and butyrate are local minima across a range of physiologically relevant pH and *E(electron donor)* values. Succinate is a key intermediate in the TCA cycle with numerous recently elucidated signalling functions^32,33^. Interestingly, succinate accumulation occurs in a number of different organisms, including bacteria such as *Escherichia coli^*34*^*, *Mycobacterium tuberculosis^*35*^*, as well as several bacterial members of the human gut microbiome^36–39^ and the bovine rumen^40,41^; fungi such as the yeast *Saccharomyces cerevisiae*^*42*^ and members of the genus *Penicillium^43^*; green algae^44^; parasitic helminths^45^; the sleeping sickness-causing parasite *Trypanosoma brucei^46^*; marine invertebrates^47^; and humans^48–51^. More specifically, our observations are consistent with the behavior of the TCA cycle under anaerobiosis and hypoxia ^52–55^. In these conditions, the reactions of the TCA cycle operate like an “incomplete fork”, with a portion of the pathway running in a reductive (‘counterclockwise’) modality, i.e. oxaloacetate sequentially reduced to malate, fumarate, and succinate. Thus, despite the fact that in these examples succinate is part of biochemical networks of higher complexity than our redox chemical space, its empirically observed accumulation is consistent with its identity as a local energy minimum. We also note that the short-chain fatty acid (SCFA) butyrate, accumulates to high (millimolar) levels in the gut lumen as a product of bacterial fermentation^56,57^. We found that this pattern - where the n-carbon linear-chain dicarboxylic acid and the fatty acid emerge as the only local minima in a large region of phase space - is conserved in the Pourbaix diagrams for redox chemical spaces with different numbers of carbon atoms (Fig. S7, S8, S12). Therefore, in analogy to succinate accumulation, it would be reasonable to search for evidence of n-carbon linear-chain dicarboxylic acid accumulation in different biological systems under physiological conditions matching the relevant region of phase space. Studying glutarate accumulation in hypoxic and/or acidic conditions would be particularly enticing, since pathways for its biosynthesis (e.g. as part of lysine metabolism) are conserved across many species.

Our results also showed that the fully reduced alkane butane (and corresponding alkanes in other redox chemical spaces) is the global minimum in the energy landscape at a wide range of pH and *E(electron donor)* values. Although bacterial alkane production has been described ^58,59^ the high volatility of these compounds likely limits their role in metabolism.

There are several caveats and limitations associated with our analysis. A first one is that our redox chemical space analysis is based solely on thermodynamics and does not account for kinetics. Thus we assume that all molecular oxidation levels are accessible, effectively ignoring kinetic constraints. A second caveat is that the set of redox transformations considered here does not account for additional constraints imposed by known enzymatic reaction mechanisms. For instance, the reduction of a hydroxycarbon (alcohol) functional group to a hydrocarbon occurs enzymatically through a C=C double-bond intermediate (for example, the reduction of malate to succinate occurs via fumarate). Therefore, a hydroxycarbon functional group that has two neighboring carbon atoms in the carbonyl oxidation level cannot undergo such a reduction using known enzymatic mechanisms. In addition, our redox chemical space ignores further biochemical details: we do not include intramolecular redox transformations (where an electron transfer within a molecule changes the oxidation level of two different carbon atoms) or keto-enol tautomerizations; we do not account for non-linear chain carbon compounds nor the different possible stereoisomers of a given molecular oxidation level (e.g. L-malate vs. D-malate) which may differ in energy; and we do not consider functional group activation chemistry (e.g. the conversion of carboxylic acids to thiols) which has an important effect on thermodynamics. Finally, our partitioning of molecules into metabolites and non-biological compounds relies on what is found in the KEGG database, which is only a proxy for the absolute set of compounds that partake in nature’s redox biochemistry.

However, despite these caveats we propose that our simplified redox chemical space is rich enough to serve as a baseline for a better understanding of the underlying thermodynamic and physicochemical principles of carbon redox biochemistry. In future work, and following recent exciting developments in the field of heuristically-aided quantum chemistry ^60–63^, our chemical space model could be expanded to include the additional types of biochemical transformations mentioned above and begin to account for kinetic accessibility. It would be particularly interesting to include carboxylation and decarboxylation reactions (both reductive/oxidative and non-reductive/non-oxidative), which would effectively connect the different n-carbon redox chemical spaces to each other, but would significantly increase the complexity of the analysis. In addition, including additional types of reactions such as aldol/retro-aldol reactions and hydrations/dehydrations would fully map the chemical space model to experimentally tractable reaction networks ^25,64–66^.

Finally, given the importance of redox chemistry at the early stages of life’s history, it is possible to think of our landscape as a generalization of the space of metabolites found in current living systems ^1,2,67^. By taking into account this extended space, future models for the rise and evolution of biochemistry^62,63,68^ could more specifically compare the evolutionary trajectory of life-as-we-know-it to alternative paths potentially involving transiently relevant molecules and reactions ^69,70^.

## Acknowledgements

We thank Arren Bar-Even for fruitful discussions and feedback, and Ron Milo, Manuel Razo-Mejia, Jennifer Wei and, Dmitrij Rappoport for valuable discussions and comments on the manuscript. The authors thank Harvard Research Computing for their support on using the Odyssey cluster. A.A.-G., A.J., and B.S.L. thank Anders G Frøseth for his generous support. J.E.G and D.S. were partially supported by grants from NASA, NSF and the Human Frontiers Science Program.

## Materials and methods

### Generation of full redox networks using RDKit

To generate the reactions, we used the RDKit cheminformatics software to design SMILES (simplified molecular-input line-entry system)^71^ reaction templates (reaction strings), which, when applied to a compound, will reduce it according to the functional groups detected. Reaction strings were created for the three redox categories of interest: reduction of carboxylic acids to aldehydes, reduction of ketones to alcohols, and reduction of alcohols to hydrocarbon. These templates are designed to be generic enough that they can be applied to any compound with the target functional group, but also with enough specificity to only generate a reaction belonging to the correct redox category.

As an illustrative example, we consider the reduction of pyruvate. Pyruvate contains two types of functional groups that can be reduced: a carboxylic acid and a ketone. The carboxylic acid can be reduced to an aldehyde, or the ketone can be reduced to a hydroxyl. To accomplish this we applied the appropriate SMILES reaction strings. The SMILES reaction string used for the ketone reduction of pyruvate to lactate is shown below. This reaction string can be visualized as a generic reduction of a ketone to a hydroxyl. The *ReactionFromSmarts* function in RDkit is used to generate a reaction object from the reaction string.

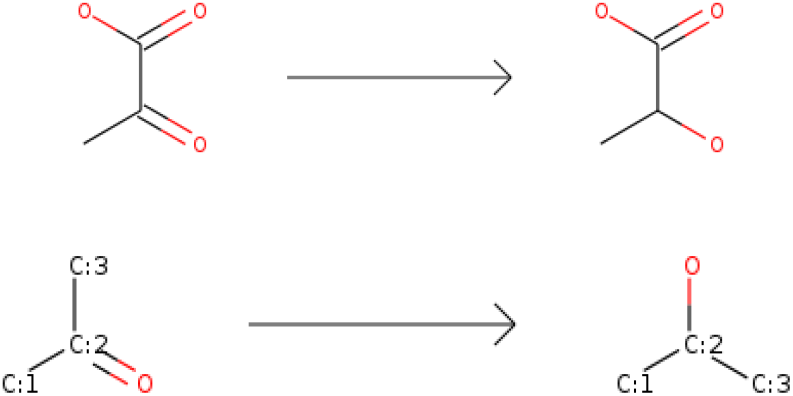

The molecular transformation encoded by the SMILES reaction string is shown above. The substrate and compound of each reaction are represented as strings and concatenated into a reaction string as follows: [#6:1][CX3:2](=O)[#6:3]≫[#6:1][CX4H1:2]([#6:3])[OX2H1]

This reaction object can be applied to any compound with a ketone functional group in order to reduce it to a hydroxyl. For cases in which the compound contains multiple target functional groups (e.g. dicarbonyls), every possible product will be generated. To generate the full network or redox reactions, these reaction strings were run iteratively, starting with the fully oxidized unbranched, carbon chain compounds of length 2 to 6 carbons. For example the seed compound for the redox chemical space of 6-carbon straight-chain molecules (i.e. the fully oxidized 6-carbon linear chain seed compound) is shown below:

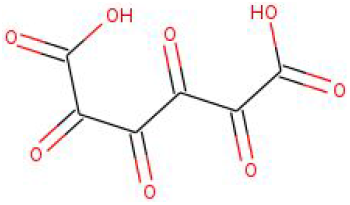

Once fully oxidized seed compound had been reduced one step at every possible carbon atom in the initial iteration, the function was repeatedly applied on the resulting products. This continued iteratively until, the fully reduced n-carbon hydrocarbon chain is obtained. Any duplicate reactions and products generated from this approach were eliminated during each iteration. Thus, a network of all possible redox reactions originating from the fully oxidized seed compound can be generated.

SMILES reaction strings

| Reaction category | Reaction strings |
| --- | --- |
| Carboxylic acids to aldehydes | [CX3:1](=O)[OX2H1]>>[CX3H1:1](=O) |
| Aldehydes to alcohols | [CX3H1:2](=O)[#6:1]>>[#6:1][CX4H2:2][OX2H1] |
| Ketones to alcohols | [#6:1][CX3:2](=O)[#6:3]>>[#6:1][CX4H1:2]([#6:3])[OX2H1] |
| Alcohols to hydrocarbons (middle) | [CX4H2:2][OX2H1]>>[CX4H3:2] |
| Alcohols to hydrocarbons (edge) | [#6:1][#6H1:2]([#6:3])[OX2H1]>>[#6:1][#6H1:2][#6:3] |

### Computing network degree distributions

The degree of a compound in the redox network is defined as the number of redox reactions - oxidations and reductions - that connect it to molecules with higher or lower oxidation level. We used the network analysis library NetworkX ^72^ in Python to compute the degree distribution of compounds in the full redox networks.

### Comparison against KEGG database

In order to classify compounds in the full redox networks as biological or non-biological, we looked for matches in the KEGG database of metabolic compounds. We did this in several steps. In order to match biological compounds against the n-carbon network, we filtered out metabolites in KEGG containing n-carbon atoms. Then, using the RDKit toolbox, we matched molecules in the networks against KEGG metabolites using their canonicalized smiles string representation ^73^. In order to additionally capture KEGG compounds that have alcohol functional groups substituted by amine or a phosphate functional groups, we visually inspected all remaining n-carbon molecules in KEGG. Finally, to capture compounds with carboxylic acids activated by Coenzyme A, we generated a list of all KEGG compounds with n-carbon atoms plus a covalently attached Co-A molecule. Manual search of this list led to the final set of biological metabolites matching compounds in our full redox networks.

### Computing the null distribution for the expected number of n-gram (single, pair and triplet) functional group patterns

Borrowing terminology from natural language processing, we call the set of all possible sequences of one, two, and three carbon functional groups the set of oxidation level n-grams. The goal is to count the number of times that each n-gram appears in the set of biological (or non biological) compounds (where N is the total number of biological compounds), and compare that against properly generated random sets of compounds (the null distribution).

### The analytical null distribution for single functional group patterns (1-grams)

We first note that a given n-gram can appear more than once in a single molecule. For example, the metabolite succinate has the functional group sequence {carboxylic acid, hydrocarbon, hydrocarbon, carboxylic acid}. Thus it contains two instances of the {carboxylic acid, hydrocarbon} 2-gram. In general, a 4-carbon linear-chain compound can have up to 4 instances of a 1-gram, up to 3 instances of a 2-gram, and up to 2 instances of a 3-gram.

Let *n*(*k*; *g*) be the number of molecules in the full redox network with k instances of 1-gram g. For example, *n*(1; *hydroxyl*) is the total number of compounds in the network with a single hydroxyl functional group. Assume a set of N molecules are randomly sampled without replacement from the network. Let *m*(*g*) be the total number of instances of the 1-gram *g* in this random set. These *m*(*g*) instances can come from different sampling configurations of molecules, each with *k* instances of the 1-gram *g*. We call *m*(*k*; *g*) be the number of molecules in the random sample with *k* instances of the 1-gram *g*.

To give a concrete example, assume a random set of size *N* = 30 molecules contains 16 instances of the n-gram g; thus *m*(*g*) = 16. One of the very many sampling configuration that can lead to this value of *m*(*g*) is sampling 17 molecules with zero instances of *g*, 10 molecules with 1 instance of *g*, and 3 molecule with two instances of *g*. Thus

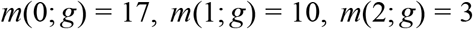

The total number of instances of the 1-gram *g* in the sample is given by:

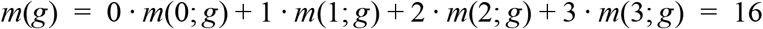

Note that the following constraint is satisfied:

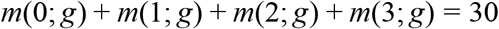

In order to compute the probability of having *m*(*g*) instances of the 1-gram *g*, we need to account for all such possible sampling configurations that add up to *m*(*g*). The number of ways of sampling *m*(*k*; *g*) molecules with *k* instances of *g* is given by 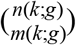. In general, given a sample size *N* and value of *m*(*g*) for n-gram *g*, the number of all possible sampling configurations that lead to that value of m(g) is given by:

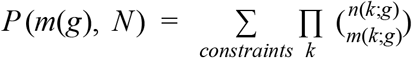

Where the summation is over terms that satisfy the following two constraints:

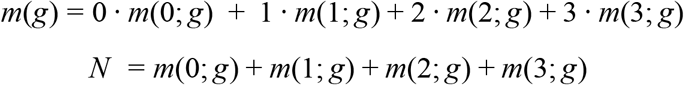

Normalizing each value of *P* (*m*(*g*), *N*) over the sum of all values leads to the probability of observing *m*(*g*) instances of the 1-gram g in a sample of size *N*, *p*(*m*(*g*), *N*). We numerically obtain the value of *n*(*k*; *g*) for *k* = 0, 1, 2, 3, 4 and g = {carboxylic acid, carbonyl, hydroxyl, and hydrocarbon}. We then numerically compute the value of *P* (*m*(*g*), *N*) by obtaining all sampling configurations that satisfy the constraints. We take *N* to be equal to the number biological compounds in the full redox network.

### The empirical null distributions for functional group pair and triplet patterns (2- and 3-grams)

Obtaining the proper null distribution for oxidation level pair and triplet patterns (2-grams and 3-grams) requires accounting for (or normalizing) for the observed single functional group statistics (1-grams). For example, the 2-gram pattern [carbonyl-carbonyl] seems to appear infrequently in the biological set of metabolites. Is this due to selection against this specific 2-gram pattern, or is it simply due to the general depletion of carbonyls (the 1-gram pattern) in the biological compounds? In order to address this, one needs to generate random sets of *N* compounds that control for or conserve the 1-gram statistics of the biological set of compounds. We numerically generate random molecules that conserve 1-gram statistics. In the case of 4-carbon linear chain molecules, we randomly choose the identity of the functional group at positions *n* = (1, 2, 3, 4) by sampling from a discrete distribution

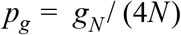

Where *g*_*N*_ is the number of instances of 1-gram *g* in the biological set, and *N* is the number of molecules in the biological set. Importantly, in order to avoid sampling carboxylic acids in the inner carbon atoms of a molecule (positions *n* = 2 *and* 3), we obtain separate functional group distributions for the inner and the outer carbon atom positions.

### Cheminformatic prediction of solubility (logS)

We used the cheminformatics software ChemAxon (Marvin 17.7.0, 2017, ChemAxon) to predict the pH-dependent solubility, logS(pH), of biological and non-biological compounds in the full redox networks. Specifically, we use the calculator plugin cxcalc logs. The cxcalc solubility calculator is based on a parametrized fragment-based model (the atom-contribution approach) fit to sets of experimental logS data ^74,75^.

### Predicting standard redox potentials with calibrated quantum chemistry approach

Our method relies on computing the electronic structure and energy of the fully protonated species of each metabolite. We obtain the smiles string for the fully protonated species and generate initial geometric conformation (with up to 10 initial conformers per metabolite) using ChemAxon (Marvin 17.7.0, 2017, ChemAxon).

All quantum chemistry calculations were performed using the Orca quantum chemistry software ^76^ version 3.0.3. We first perform a geometry optimization using density functional theory with the B3LYP functional ^77^, with Orca’s DefBas-2 basis set, COSMO implicit solvation ^78^, and D3 dispersion correction ^79^. We then perform an additional electronic single point energy (SPE) using the double-hybrid functional B2PLYP ^19,20^ (with the DefBas-5 Orca basis set, COSMO implicit solvation ^78^, and D3 dispersion correction ^79^). We note that the model chemistry selected - the combination of DFT functional, basis set, implicit solvent model, and dispersion correction for both the geometry optimization and the single point energy - was done based on a combinatorial exploration of different options.

We Boltzmann average the electronic energies of compounds, and obtain the difference in electronic energies of products and substrates for all redox reactions in the full redox networks. Every redox reaction (in the direction of reduction) was balanced by a hydrogen molecule H_2_ in the substrate side of the equation. Reductions of carboxylic acids to aldehydes and reductions of alcohols to hydrocarbons were balanced with a water molecule H_2_O in the product side of the equation.

The difference in product and substrate electronic energies is an estimate of the chemical redox potential for the fully protonated species, E^o^(fully protonated species). In order to convert this chemical potential to the biochemical potential at pH = 7, E^o^’(pH=7), we use pKa estimates from Chemaxon (Marvin 17.7.0, 2017, ChemAxon) and the Alberty Legendre transform.

Our approach relies on several approximations, such as ignoring vibrational enthalpy and entropy contributions to the formation Gibbs energy of compounds. In order to correct for systematic in the quantum chemistry methodology and the empirical pKa estimates used, we calibrate predictions against available experimental data using linear regression.

### Predicting standard redox potentials with the group contribution method

The group contribution method relies on a fragment-based decomposition of compounds into group, each of which is assigned a group energy based on available experimental data ^15–18^. Reaction energy estimates are obtained by taking the difference of the group energy vectors of products and substrates. We used the group contribution method as implemented by Noor et al. ^18^ to estimate the redox potentials of the set of linear-chain carbon redox reactions with experimental values.

### Determining statistical significance

For all tests of statistical significance (i.e. differences in solubilities, n-gram counts, octanol-water partition coefficients, Gibbs energies of biological vs. non-biological compounds) we performed Welch’s unequal variance t-test, which is an adaptation of Student’s t-test that does not assume equal variance.

