## Supplementary Figures for "A thermodynamic atlas of carbon redox chemical space"

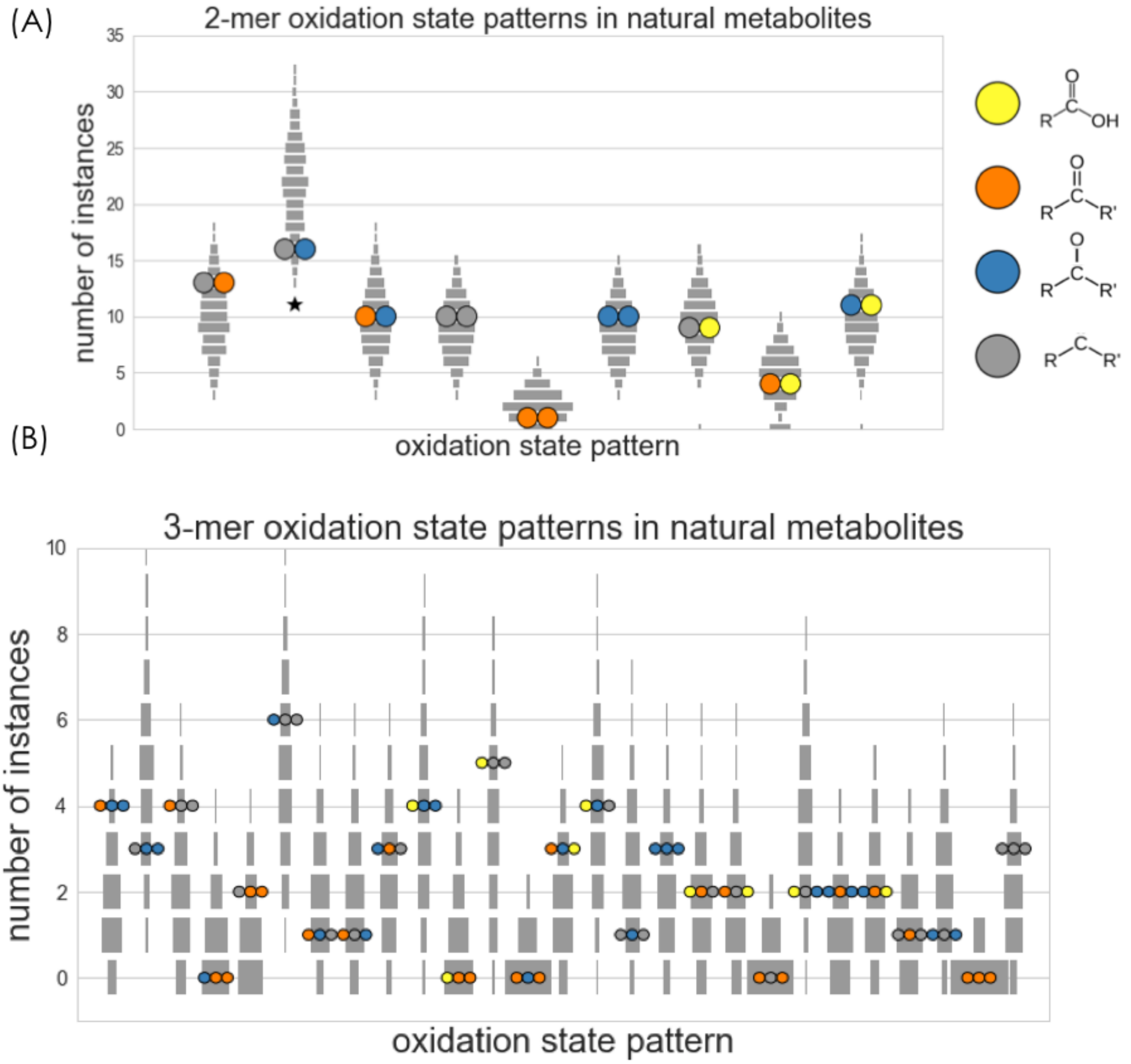


**Figure S1: Enrichment and depletion of functional group pair and triplet patterns in 4-carbon linear-chain redox chemical space.** A) The number of times each possible pattern of nearest neighbor functional group pairs appears in the set of biological metabolites are shown as pairs of colored circles. Gray squares correspond to the empirically-derived null distributions for randomly sampled sets of molecules from the network. The null distributions account for the single functional group (1-mer) statistics (see Methods). The pattern *hydrocarbon-alcohol* is depleted in the biological compounds, but with weak statistical significance (p = 0.05). B) The number of times each possible pattern of functional group triplets appears in the set of biological metabolites. No patterns are significantly enriched or depleted in the set of biological metabolites.

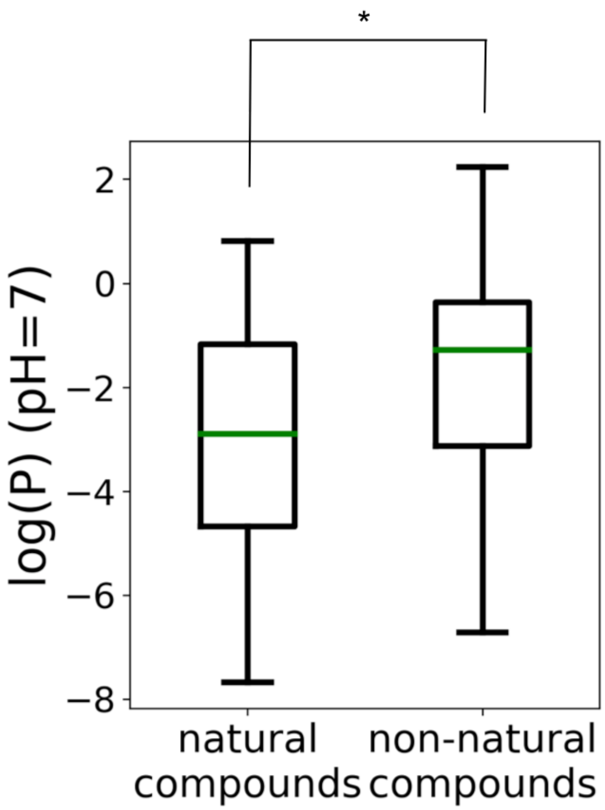


**Figure S2: Octanol water partition coefficients of biological vs. non-biological compounds in 4-carbon linear-chain redox chemical space.** Comparison of predicted octanol-water partition coefficient log(P) at pH=7 for biological and non-biological compounds in the 4-carbon linear-chain redox network. This is also known as the distribution coefficient (logD). biological compounds have significantly lower logP(pH=7) than the non-biological set ($p < 0.01$).

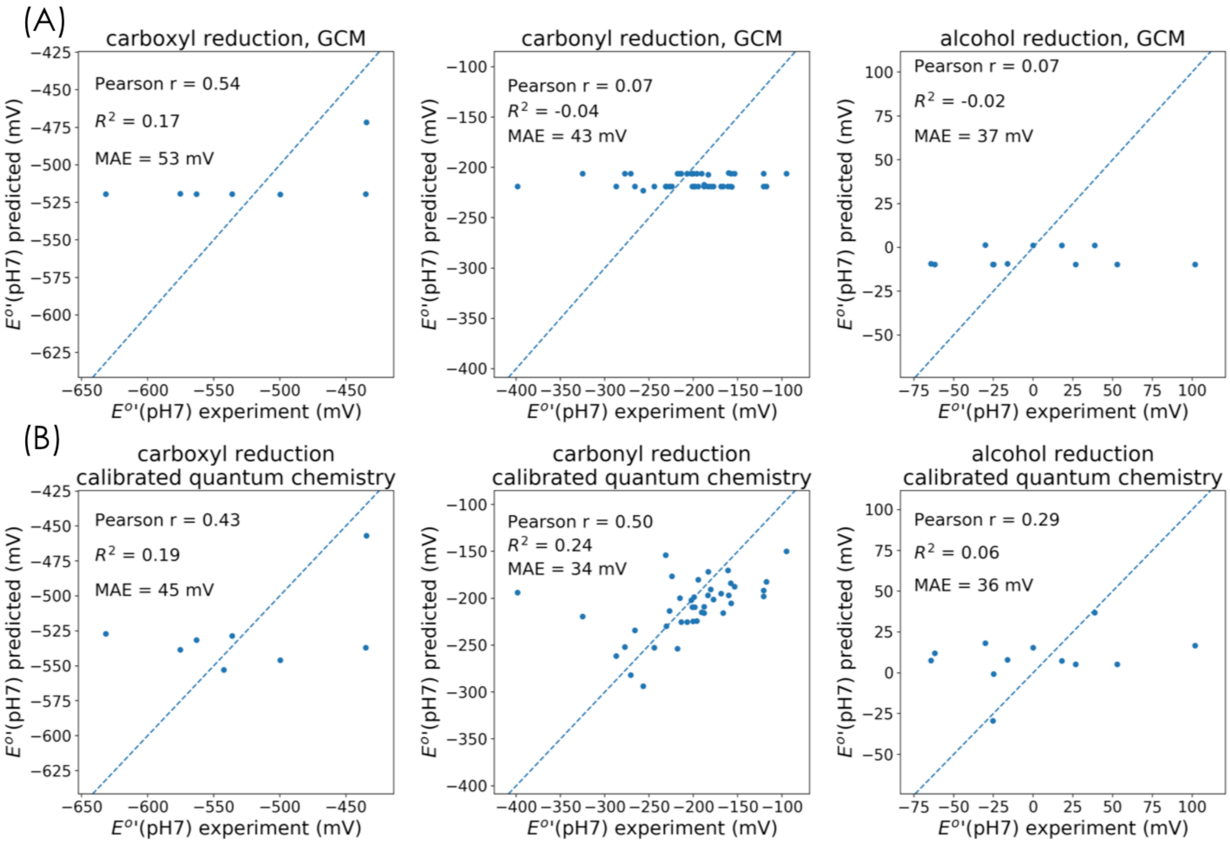


**Figure S3: Accuracy of group contribution method and calibrated quantum chemistry redox potential predictions.** Experimental data was obtained from the NIST database for Thermodynamics of Enzyme-Catalyzed Reactions (TECRDB) A) Group contribution method prediction accuracies for reduction potentials of carboxylic acids, carbonyls, and alcohol functional groups in linear chain compounds. A) Calibrated quantum chemistry prediction accuracies for reduction potentials of carboxylic acids, carbonyls, and alcohol functional groups in linear chain compounds. Quantum chemistry calculations were performed using density functional theory with a double hybrid functional (B2PLYP), and calibrated against experimental data using linear regression.

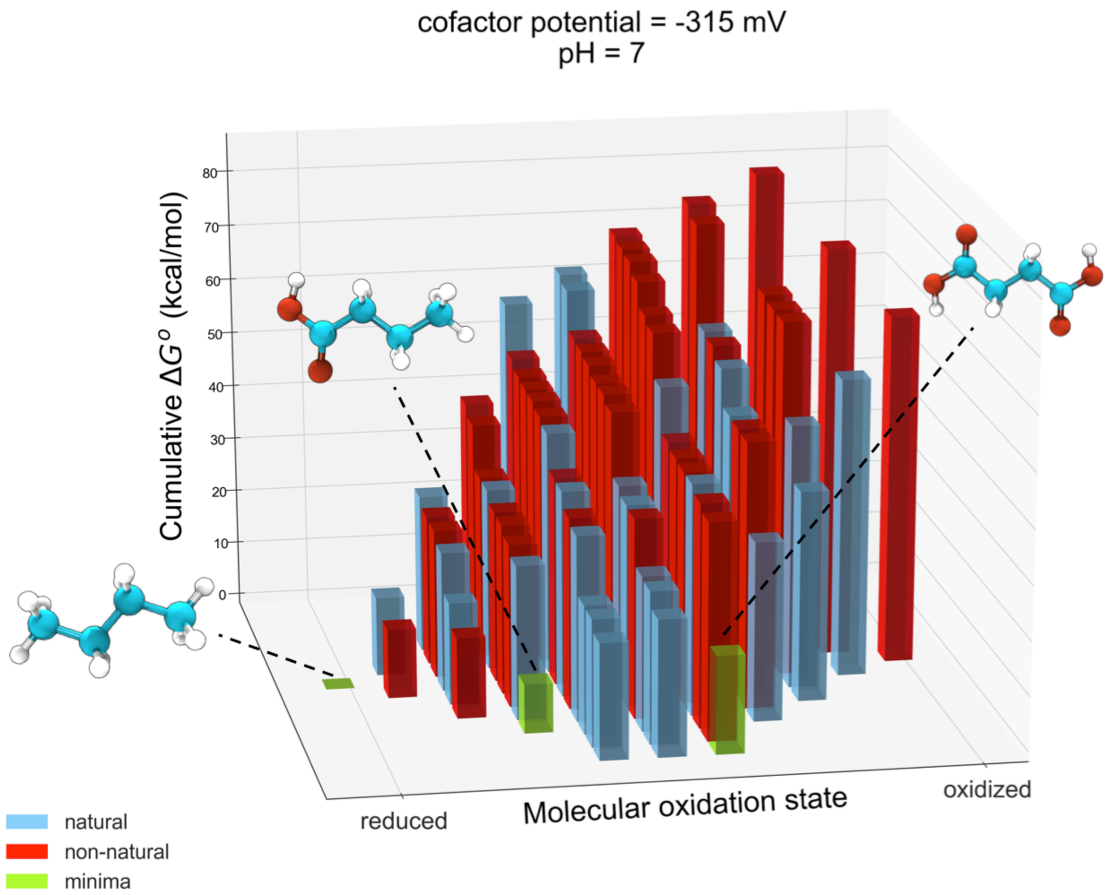


**Figure S4: Thermodynamic landscape of 4-carbon linear chain redox network at pH=7 and cofactor potential E(cofactor) = -315 mV.** Gibbs energies are normalized relative to the metabolite with the lowest energy (butane). Thus the cumulative Gibbs energies of a metabolite is obtained by summing up the Gibbs reaction energies of all reactions leading to it from the reference metabolite. Compounds within a column (i.e. with the same molecular oxidation state) are sorted according to their energies. The three compounds - butane, butanoic acid, and succinate - which are local minima in the thermodynamic landscape are shown. These local minima have lower energy than any of their neighboring molecules which are accessible by either a reduction or an oxidation.

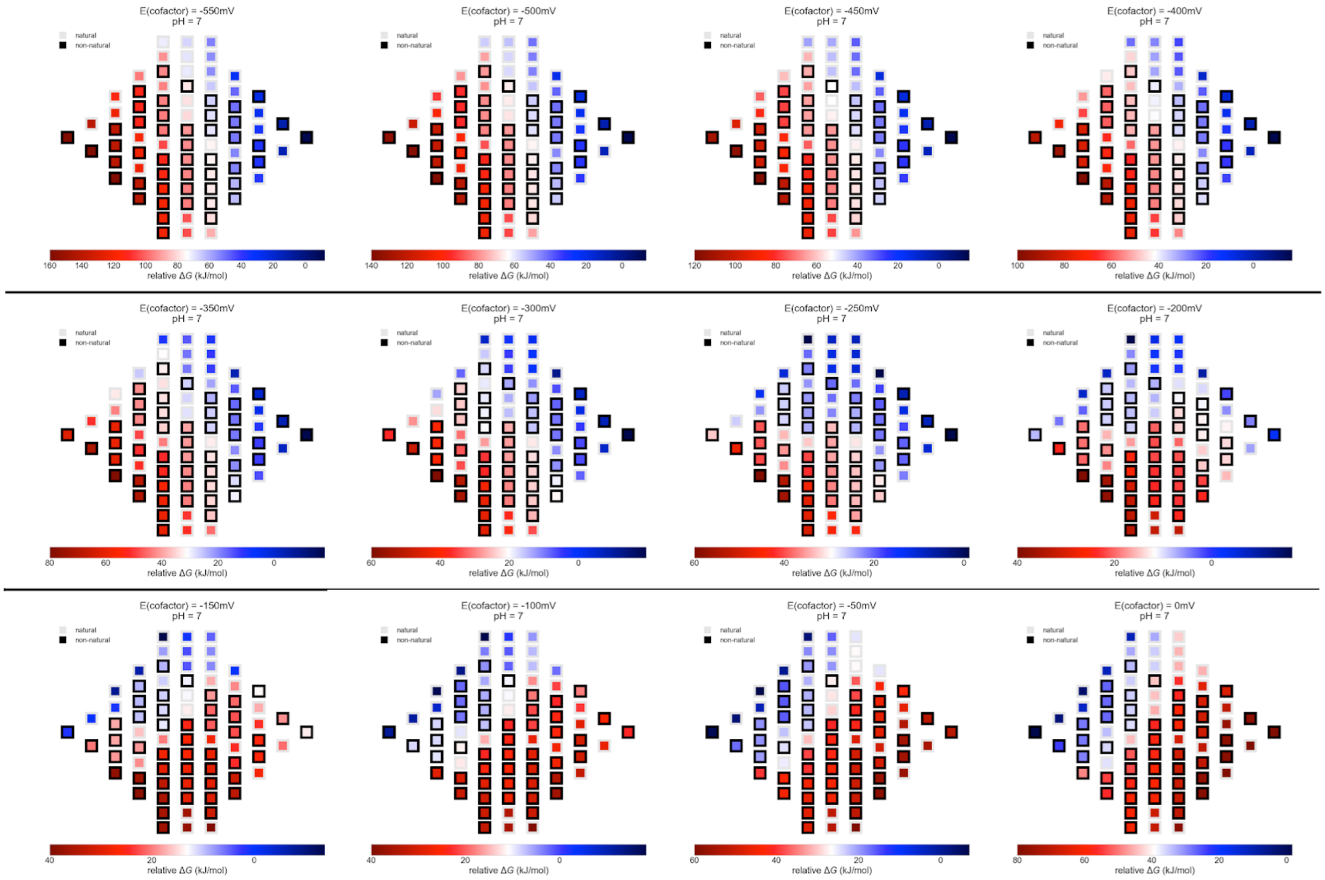


**Figure S5: Thermodynamic landscape of the 4-carbon network at a fixed value of pH and a range of values of electron donor potential (-550 mV to 0 mV).** Relative Gibbs energies of compounds is color-coded with blue (low energies) to red (high energies). Higher values of the electron donor potential energetically drive the redox chemical space towards more oxidized compounds, while lower values energetically drive the redox chemical space towards more reduced compounds.

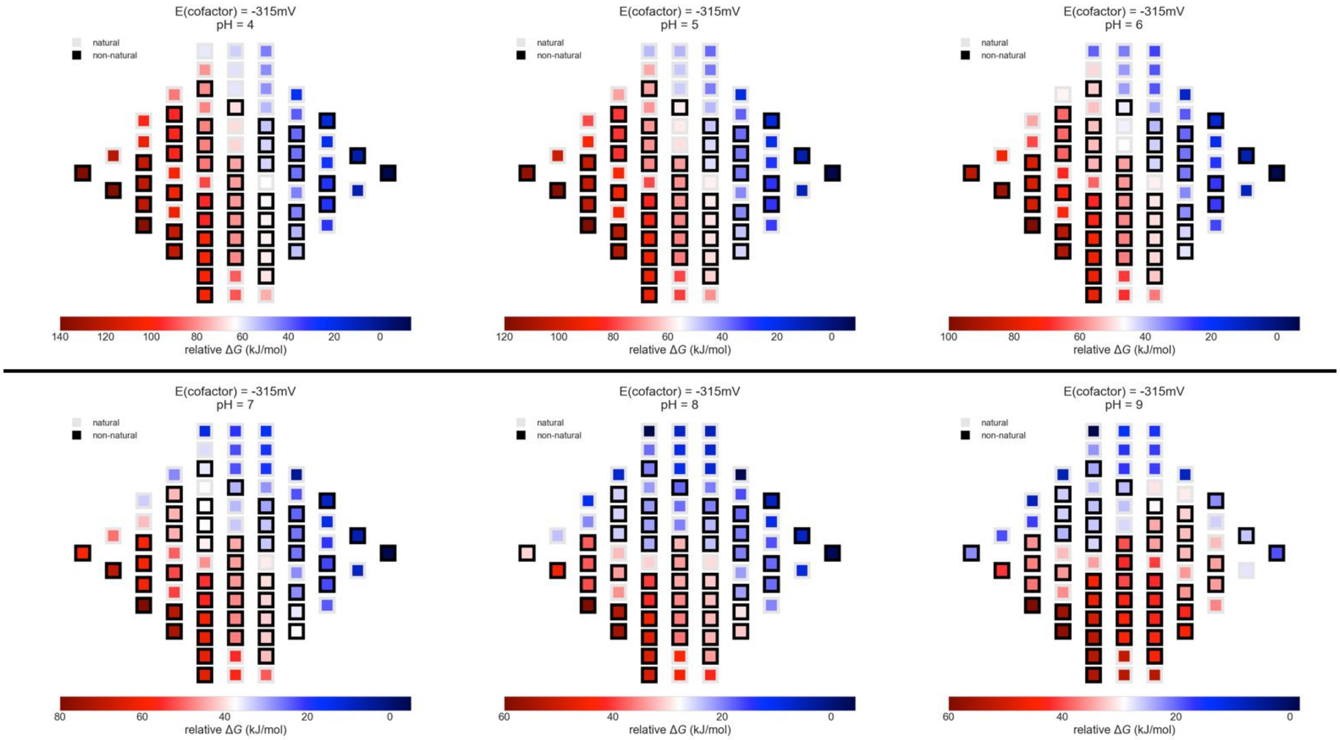


**Figure S6: Thermodynamic landscape of the 4-carbon network at a fixed value of electron donor potential and a range of values of pH (4-9).** Relative Gibbs energies of compounds is color-coded with blue (low energies) to red (high energies). Acidic pH drives the landscape towards more reduced compounds, while basic pH drives the landscape to more oxidized compounds.

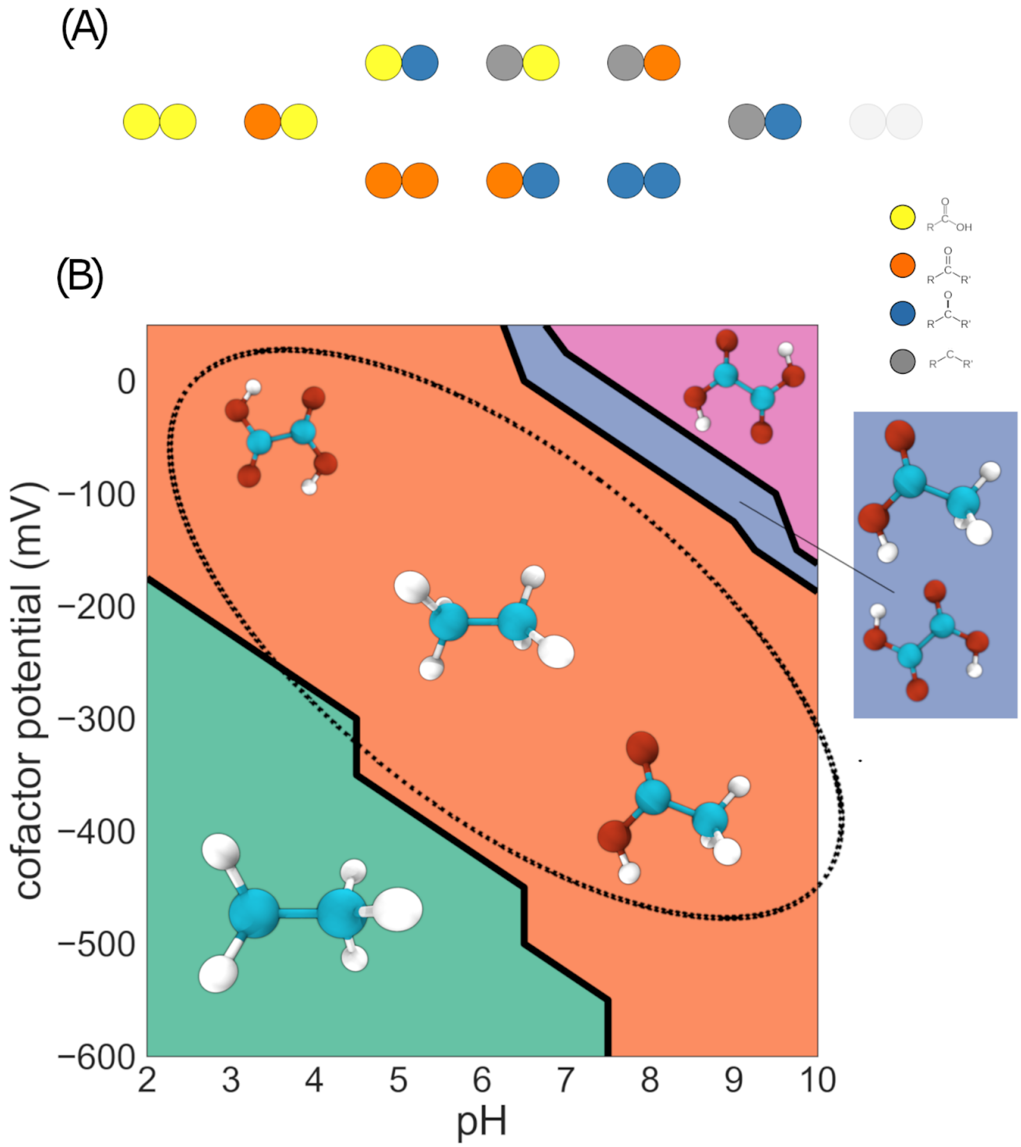


**Figure S7: The 2-carbon redox chemical space.** A) The subset of molecules in 2-carbon linear-chain redox chemical space that match biological metabolites in the KEGG database. Carbon atoms are represented as colored circles, with each color corresponding to an oxidation state: yellow = carboxylic acid; orange = carbonyl; blue = hydroxycarbon; gray = hydrocarbon. Only the fully reduced hydrocarbon ethane does not match a biological metabolite. B) Pourbaix phase diagram for the 2-carbon linear chain redox chemical space. Molecules that are local minima in the energy landscape at each region of pH vs. E(electron donor/acceptor) phase space are shown. At low pH and E(electron donor/acceptor) values, ethane is both the global and the only local minimum energy compound, while at high pH and E(electron donor/acceptor) values, the fully oxidized oxalate is both the global and the only local minimum energy compound. The dashed circle highlights the region of phase space where the dicarboxylic acid oxalate and the 2-carbon fatty-acid acetate (along with ethane) are the local minima.

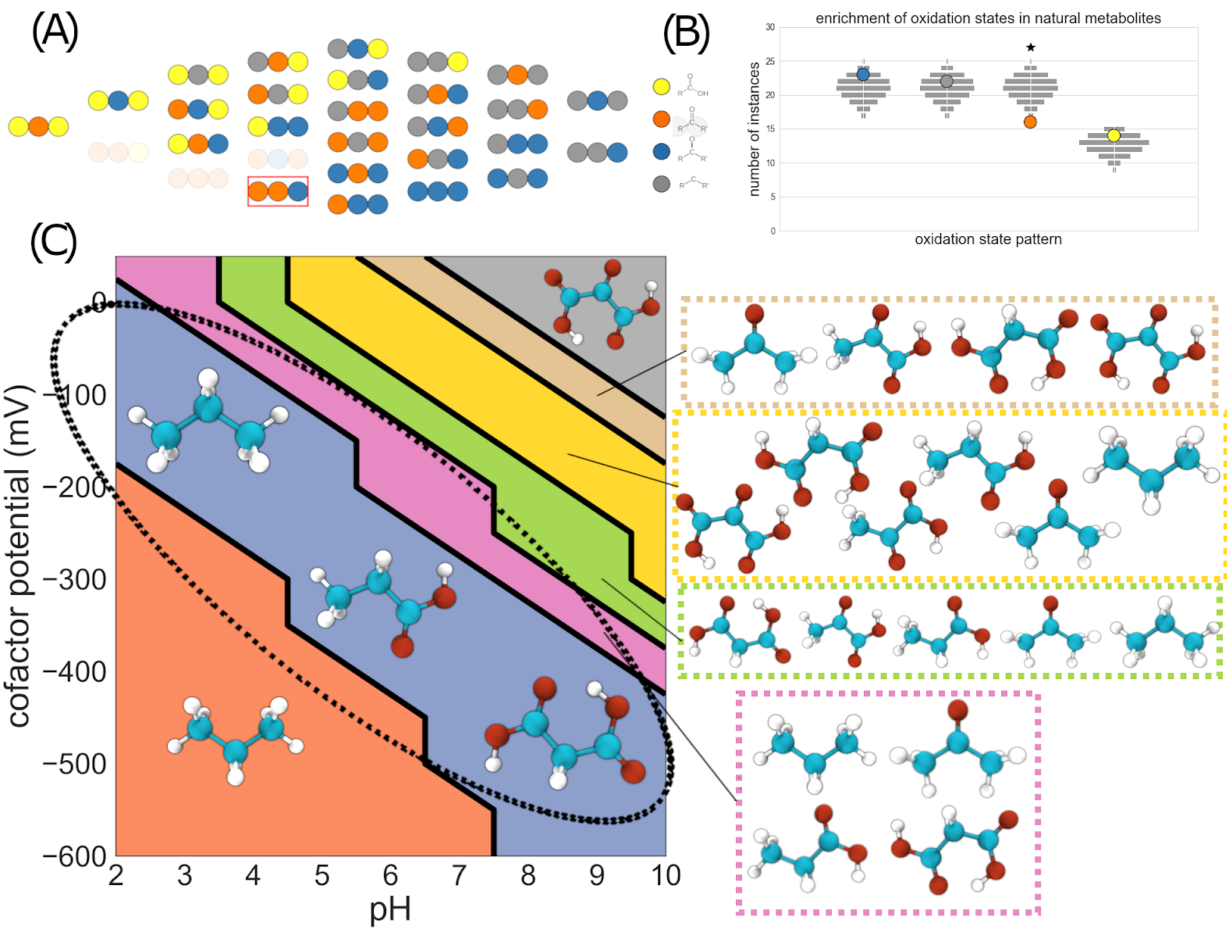


**Figure S8: The 3-carbon redox chemical space.** A) The subset of molecules in 3-carbon linear-chain redox chemical space that match biological metabolites in the KEGG database. Carbon atoms are represented as colored circles, with each color corresponding to an oxidation state: yellow = carboxylic acid; orange = carbonyl; blue = hydroxycarbon; gray = hydrocarbon. B) Enrichment and depletion of functional groups in the set of biological compounds. The vertical position of each colored circle corresponds to the number of times each functional group appears in the set of biological compounds. The light gray squares show the corresponding expected null distributions for random sets of molecules sampled from redox chemical space. C) Pourbaix phase diagram for the 3-carbon linear chain redox chemical space. Molecules that are local minima in the energy landscape at each region of pH vs. E(electron donor/acceptor) phase space are shown. At low pH and E(electron donor/acceptor) values, propane is both the global and the only local minimum energy compound, while at high pH and E(electron donor/acceptor) values, the fully oxidized 3-carbon compound (oxomalonate) is both the global and the only local minimum energy compound. The dashed circle highlights the region of phase space where the dicarboxylic acid malonate and the 3-carbon fatty-acid propionate (along with propane) are the only local minima.

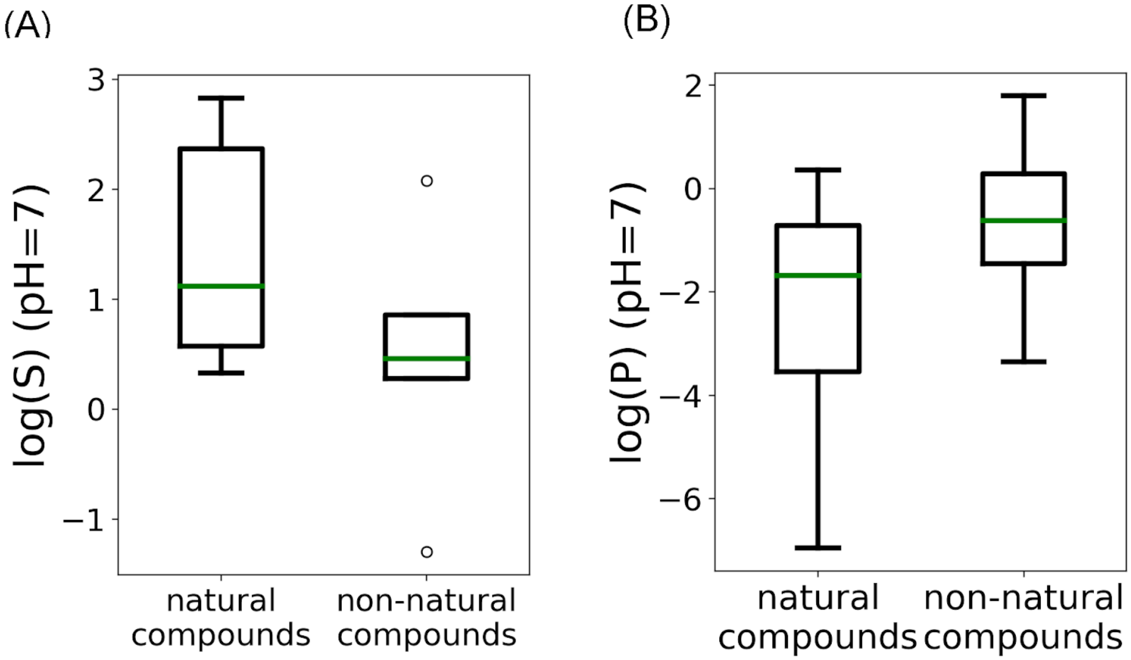


**Figure S9: Solubility and octanol-water partition coefficients of biological and non-biological compounds in 3-carbon redox chemical space.** Comparison of predicted aqueous solubility log(S) at pH=7 for biological and non-biological compounds in the 3-carbon linear-chain redox chemical space. Although the biological compounds in the 3-carbon redox chemical space tend to have higher solubilities and lower octanol-water partition coefficients at pH=7, the differences are not statistically significant (Welch's t-test).

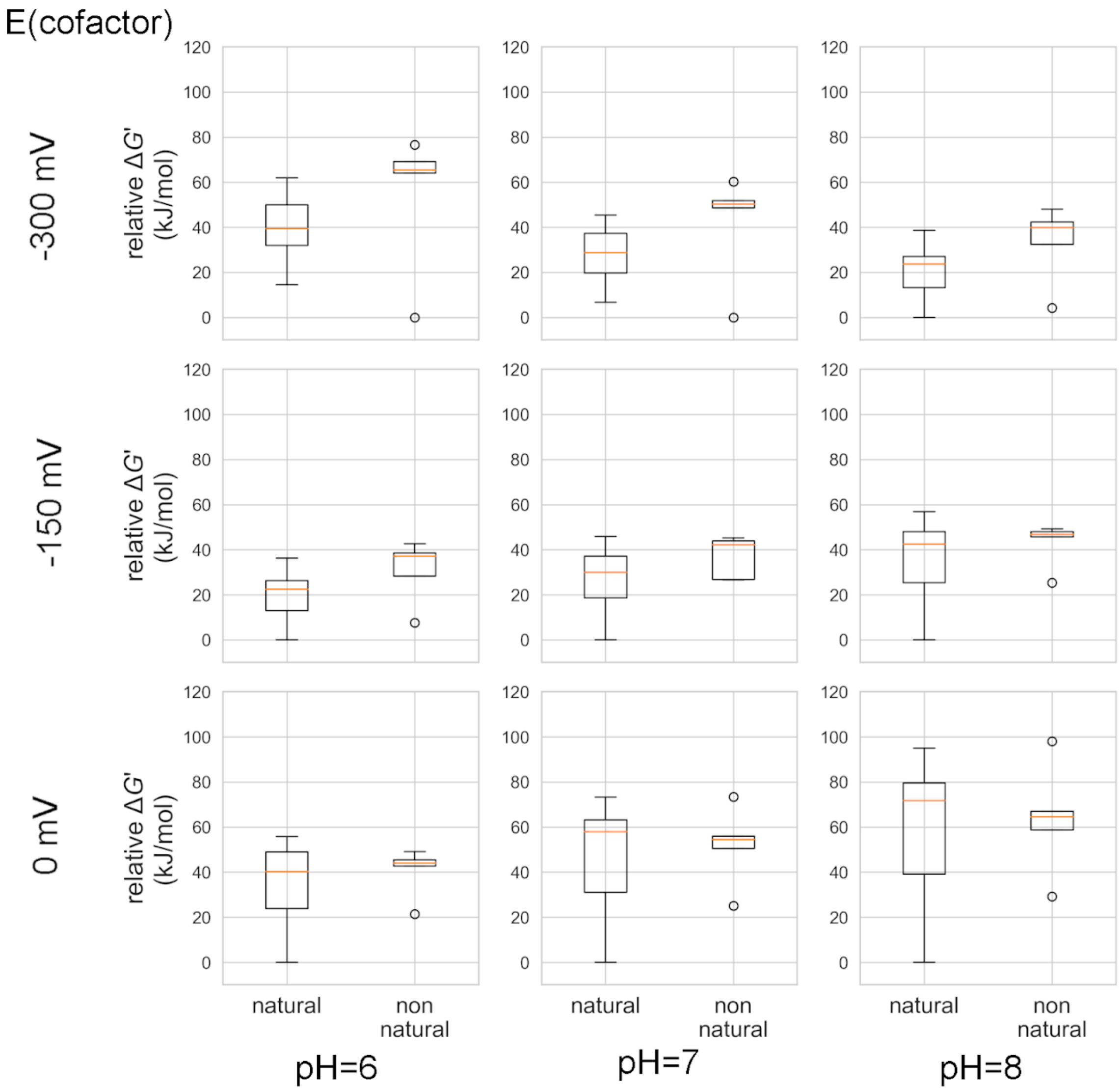


**Figure S10: Relative energies of biological and non-biological compounds in 3-carbon redox chemical space.** Relative Gibbs energies of biological and non-biological compounds in the 3-carbon redox chemical space for a range of pH and E(electron donor/acceptor) values. At each value of pH and E(electron donor/acceptor), Gibbs energies are normalized relative to the compound with the lowest energy. Although biological metabolites tend to have, on average, lower energies than the non-biological compounds, the differences are not statistically significant (Welch's t-test, $p>0.05$). The low energy of the fully reduced propane across conditions tends to bring down the average relative energy of the non-biological compounds.

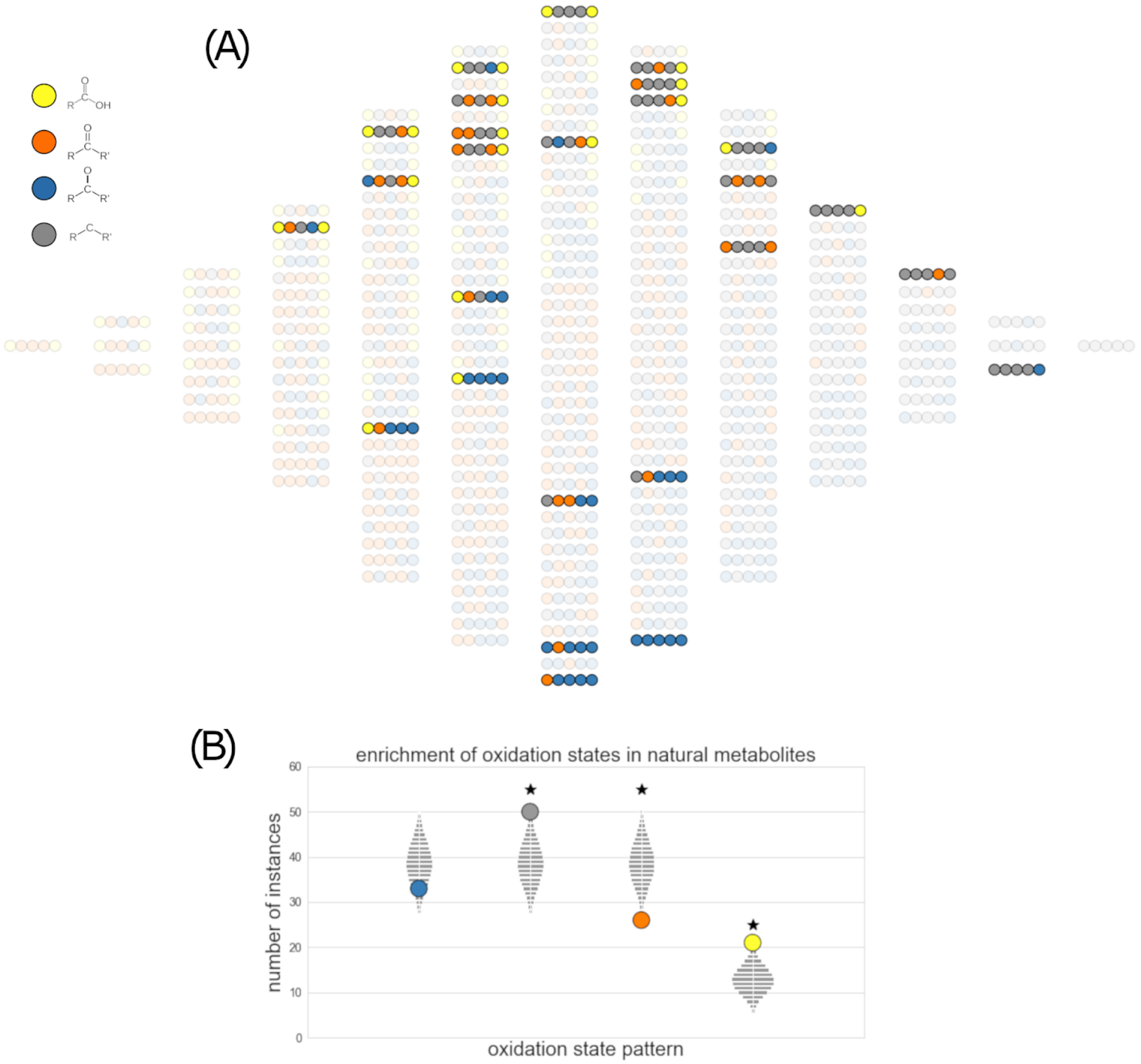


**Figure S11: The 5-carbon linear-chain redox chemical space.** A) The subset of molecules in 5-carbon linear-chain redox chemical space that match biological metabolites in the KEGG database. Carbon atoms are represented as colored circles, with each color corresponding to an oxidation state: yellow = carboxylic acid; orange = carbonyl; blue = hydroxycarbon; gray = hydrocarbon. B) Enrichment and depletion of functional groups in the set of biological compounds. The vertical position of each colored circle corresponds to the number of times each functional group appears in the set of biological compounds. The light gray squares show the corresponding expected null distributions for random sets of molecules sampled from redox chemical space.

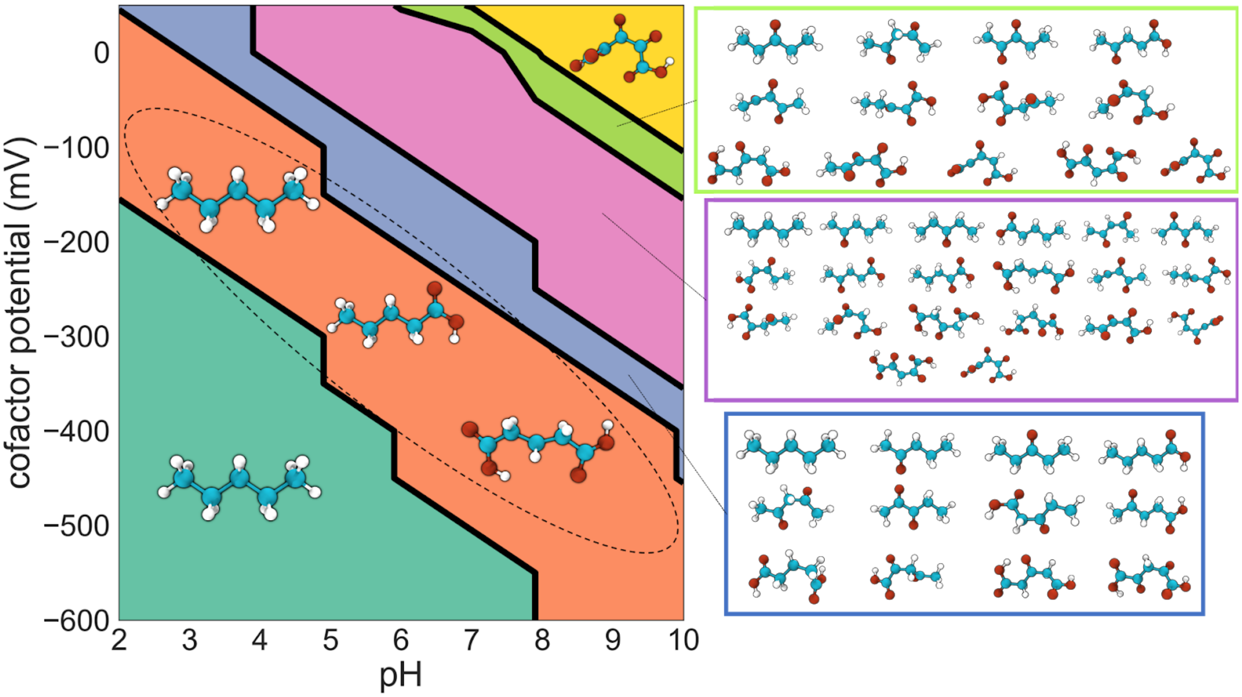


**Figure S12: Pourbaix phase diagram for the 5-carbon linear chain redox chemical space.** Molecules that are local minima in the energy landscape at each region of pH vs. E(electron donor/acceptor) phase space are shown. At low pH and E(electron donor/acceptor) values, pentane is both the global and the only local minimum energy compound, while at high pH and E(electron donor/acceptor) values, the fully oxidized 5-carbon compound (2,3,4-trioxoglutarate) is both the global and the only local minimum energy compound. The dashed circle highlights the region of phase space where the dicarboxylic acid glutarate and the 5-carbon fatty-acid valerate (along with pentane) are the only local minima.

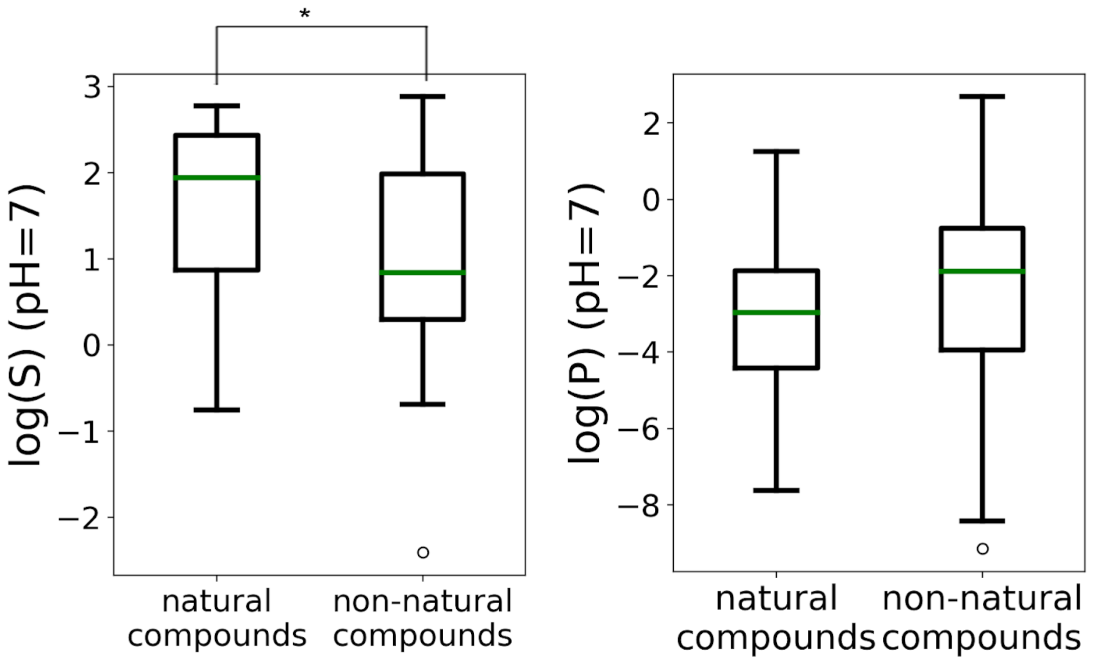


**Figure S13: Solubility and octanol-water partition coefficients of biological and non-biological compounds in 5-carbon linear-chain redox chemical space.** Comparison of predicted aqueous solubility log(S) and predicted octanol-water partition coefficient log(P) at pH=7 for biological and non-biological compounds in the 5-carbon linear-chain redox chemical space. biological compounds have statistically significantly higher solubilities (Welch's t-test) than the non-biological set ($p < 0.05$).

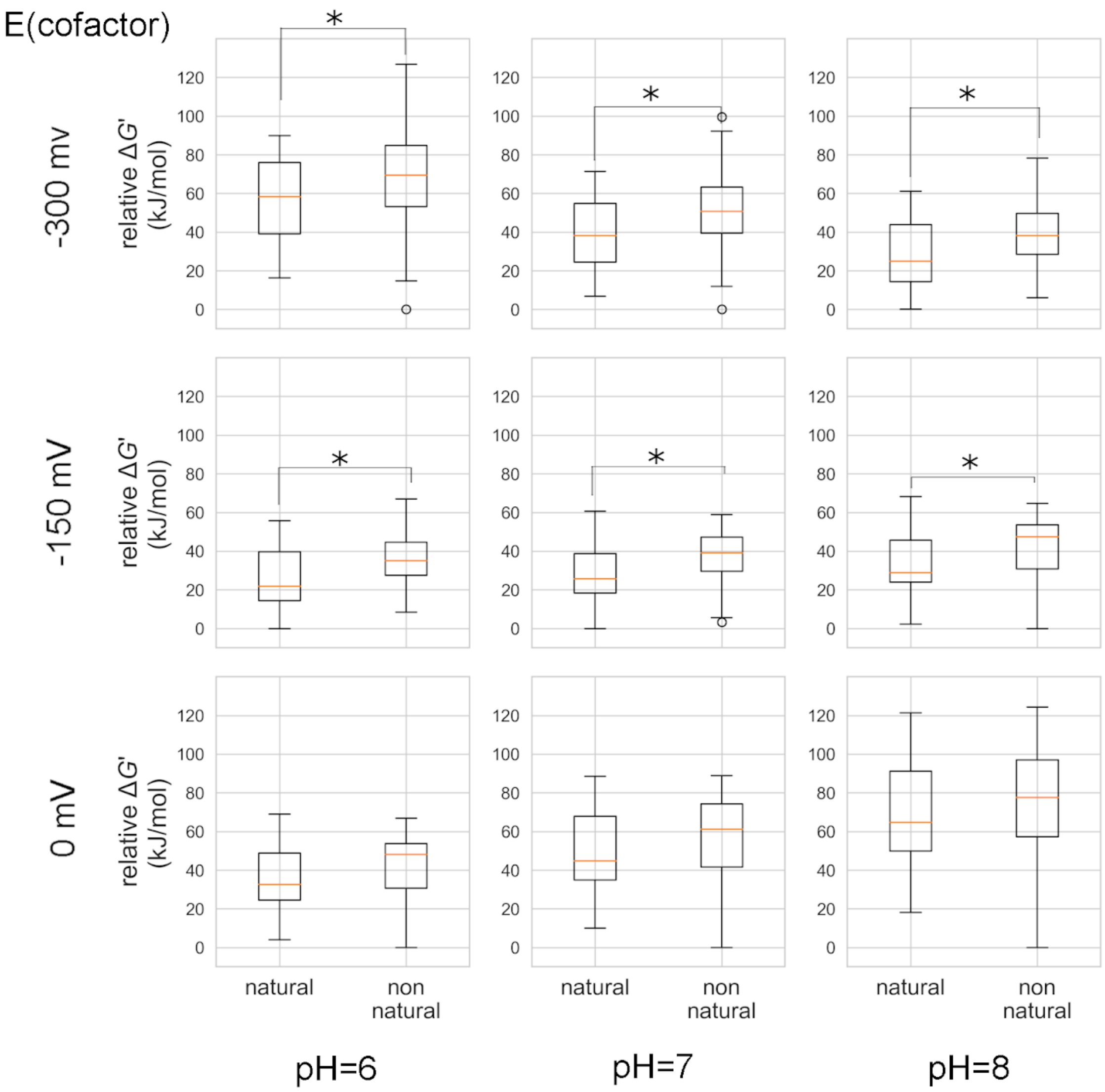


**Figure S14: Relative energies of biological and non-biological compounds in 5-carbon linear-chain redox chemical space.** Relative Gibbs energies of biological and non-biological compounds in the 5-carbon redox chemical space for a range of pH and E(electron donor/acceptor) values. At each value of pH and E(electron donor/acceptor), Gibbs energies are normalized relative to the compound with the lowest energy. An asterisk indicates a statistically significant difference in the average values (Welch's t-test) ($p<0.05$).
